# The conventional dendritic cell 1 subset primes CD8^+^ T cells and traffics tumor antigen to drive anti-tumor immunity in the brain

**DOI:** 10.1101/2021.09.13.460088

**Authors:** Jay A. Bowman-Kirigin, Brian T. Saunders, Rupen Desai, Anthony Z. Wang, Max O. Schaettler, Connor J. Liu, Alexandra J. Livingstone, Dale K. Kobayashi, Vivek Durai, Nicole M. Kretzer, Gregory J. Zipfel, Eric C. Leuthardt, Joshua W. Osbun, Michael R. Chicoine, Albert H. Kim, Kenneth M. Murphy, Tanner M. Johanns, Bernd H. Zinselmeyer, Gavin P. Dunn

## Abstract

The central nervous system (CNS) antigen presenting cell (APC) which primes anti-tumor CD8^+^ T cell responses remains undefined. Elsewhere, the conventional dendritic cell 1 (cDC1) performs this role. However, steady-state brain cDC1 are rare; cDC localize to choroid plexus and dura. Using preclinical glioblastoma models and cDC1-deficient mice, we explored the role of cDC1 in CNS anti-tumor immunity. We determined that cDC1 mediate checkpoint blockade-induced survival benefit and prime neoantigen-specific CD8^+^ T cells against brain tumors. We observed that cDC, including cDC1, isolated from the tumor, the dura, and the CNS-draining cervical lymph nodes harbored a traceable fluorescent tumor-antigen. In patient samples, we observed several APC subsets (including the CD141^+^ cDC1-equivalent) infiltrating glioblastomas, meningiomas, and dura. In these same subsets, we identified a tumor-specific fluorescent metabolite of 5- aminolevulinic acid, which labels tumor cells during fluorescence-guided glioblastoma resection. Together, these data elucidate the specialized behavior of cDC1 and suggest cDC1 play a significant role in CNS anti-tumor immunity.

**One Sentence Summary:** CNS cDC1 engage in previously undefined behavior to establish immune responses against brain tumors.

## INTRODUCTION

Glioblastoma (GBM) is the most common CNS primary malignancy and remains inexorably lethal. The standard of care involving surgery, chemotherapy, and radiation treatment leads to a median survival of only 15-20 months (*1–3*). As immunotherapy has changed the care of many cancer types, there has been significant interest in applying immune-based treatments to patients with GBM, especially as there is a growing understanding that the CNS is not as immunoprivileged as long perceived (*4, 5*). However, despite numerous clinical trials, there remain no FDA-approved immunotherapies for GBM (*6–8*).

Several critical issues underlie the difficulty of harnessing immune-potentiating therapies to treat GBM. First, GBM represents a severe example of cancer immunoediting in patients (*9*) and exhibits substantial evidence of immunosuppression. A range of immunologic deficits has been described in GBM patients including lymphopenia (*10, 11*), intrinsic immune suppression by tumor cells (*12–14*) and overexpression of checkpoint molecules such as PD-L1 (*15, 16*), among others (*17–19*). These observations have led many to describe this cancer as “cold.” Secondly, because GBM tumors are molecularly heterogeneous and are comprised of at least several subclonal populations (*20, 21*), T cell antigen targets are also variably distributed in the tumor mass and not uniformly expressed.

Our incomplete understanding of the cellular basis for the development of CNS immune responses also remains a significant barrier to successful immunotherapy in the brain. Specifically, the mechanistic underpinnings of antigen presentation leading to T cell priming of brain tumor specific T cells remains undefined. It has been appreciated that, unlike other sites in the body, the brain parenchyma does not harbor dendritic cells in the resting state (*22–24*) and is not vested with classical lymphatic and secondary lymphoid structures (*22, 23, 25-27*). However, the recent characterization of lymphatic channels within the dura mater of the cerebral meninges has provided new insights into how antigen could encounter APC and be trafficked to draining lymph nodes (*28–31*). Although these foundational observations highlight the importance of the anatomic basis for lymphatic drainage in the CNS, the cellular basis for the connection of this system to antigen presentation remains unclear. The existence of cell-migration from the CNS to cervical lymph nodes has been reported (*32–35*), but this work was largely restricted to exogenously derived cultured monocyte-derived DC. Thus, there remains a lack of understanding regarding the cellular basis for endogenous antigen presentation in CNS anti-tumor immunity.

Herein, we characterized the role of the cDC1 subset during the endogenous immune response to brain tumors. The cDC1 phagocytizes antigen, migrates to lymph nodes, and cross-presents antigen to prime CD8 T cell responses. While this subset is required for effective immune responses to a range of tumor types (*36–43*), it is unclear whether these cells perform a similarly obligate role in the immune response to CNS tumors. In this study, we demonstrate that cDC1 are required both to mount neoantigen-specific immune responses and to respond to checkpoint blockade in wild-type syngeneic orthotopic preclinical models of GBM. We also observed tumor-antigen containing cDC1 within the tumor, the dura, and in the cervical lymph nodes. We extended these findings to patients, and observed the infiltration of cDC (including the analogous cDC1 subset) in human tumors as well as the dura. Finally, from surgical cases in which 5-aminolevulinic acid (5-ALA) was employed to facilitate fluorescence guided GBM resection (*44–47*), we identified the presence of its metabolite PPIX (*48, 49*), within tumor infiltrating cDCs and monocytes, but not in tumor infiltrating T cells nor in matched peripheral APC counterparts. Together, these findings clarify the unique behavior of CNS cDC1 and demonstrate that cDC1 play a critical role in the CNS anti-tumor immune response.

## RESULTS

### cDC1 are recruited to the CNS tumor microenvironment and mediate protection by checkpoint blockade

We first investigated whether cDC are recruited to the brain tumor microenvironment in orthotopic, syngeneic murine GBM. We transplanted the C57BL/6-derived GL261 tumor into the cerebral hemispheres of wild type mice and identified that all dendritic cell subsets (defined as CD45^+^, F4/80^-^, I-Ab^+^, CD11c^+^), including cDC1 (additionally XCR1^+^, SIRPα^-^, Ly-6C^-^), infiltrate brain tumors in much greater numbers than in sham controls (Figure 1A). Similarly, we observed the presence of greater numbers of DC subsets within orthotopically transplanted C57BL/6-derived CT2A tumors compared to sham controls. To visualize cDC1 in brain tumors, we employed a cDC1 reporter mouse to monitor cDC1 by flow cytometry and imaging. In the *Snx22^GFP/+^* knock-in mouse, cDC1 specifically and constitutively express GFP (*50*). When GL261 or CT2A cells were transplanted into the brains of *Snx22^GFP/+^* mice, cDC1 and cDC2 subsets infiltrated brain tumors by flow cytometry, and GFP expression was restricted to the XCR1^+^ cDC1 subset (Figure 1B). To determine the location of cDC1 within GL261 tumors, we used 2-photon microscopy to identify GFP expressing cDC1. Whereas cDC1 were vanishingly scarce within the brain parenchyma in the steady state, cDC1 infiltrated the tumor parenchyma in extravascular spaces in *Snx22^GFP/+^* mice (Figure 1C). These data demonstrate that DC, including cDC1, infiltrate orthotopically transplanted GBM.

**Figure 1.**
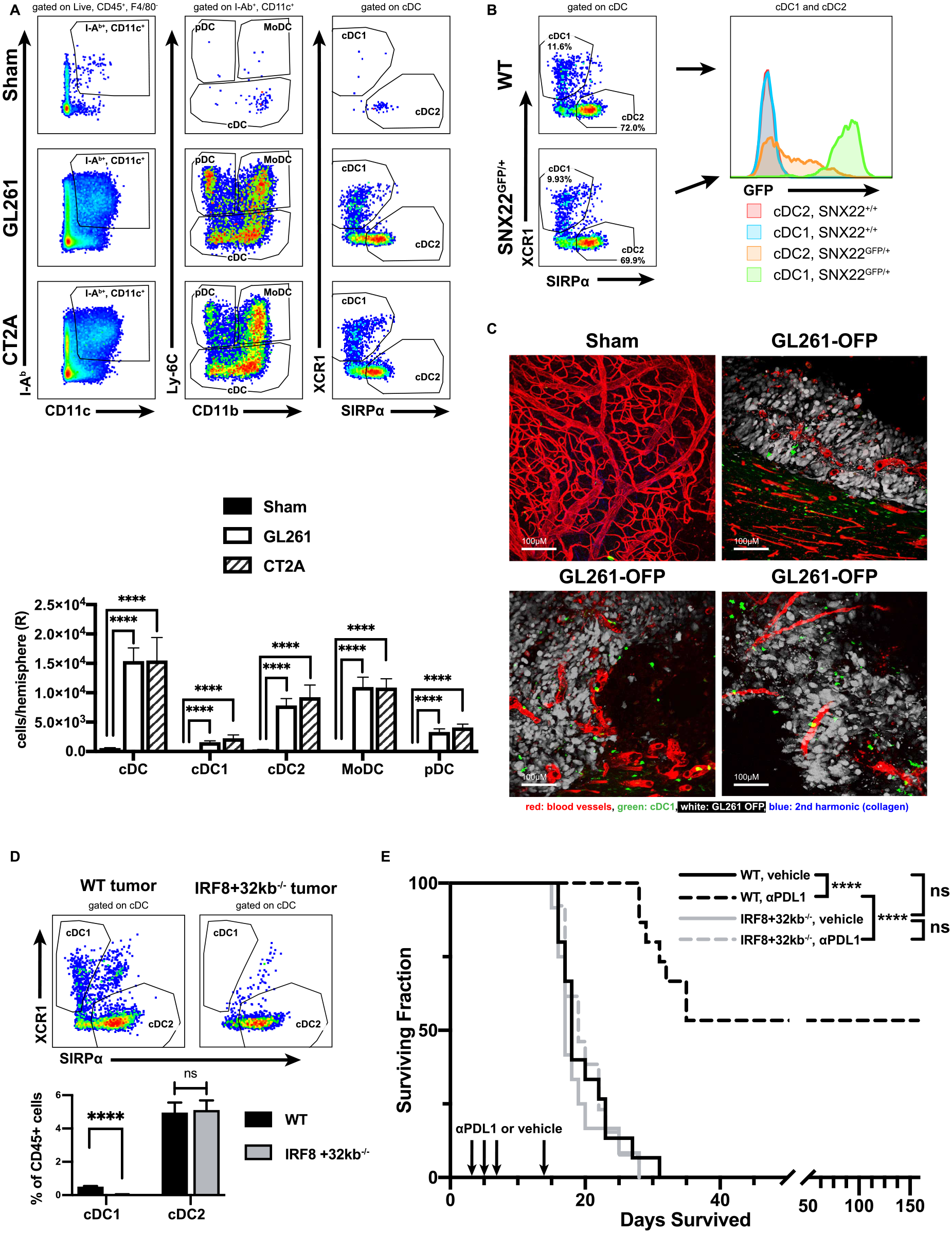
cDC1 infiltrate mouse glioma, and are required to mediate benefit from αPD-L1 therapy. **(A)** cDC1, cDC2, MoDC and pDC infiltration of brains of sham treated vs. GL261 or CT2A intracranially injected mice assessed by flow cytometry at day 14 following tumor induction. **(B)** cDC1/cDC2 profile and GFP expression of *SNX22^GFP/+^* mouse. **(C)** 2-photon microscopy of sham injected brain, or GL261-OFP injected brain at various regions in tumor. **(D)** WT vs *IRF8+32kb^-/-^* GL261 brain tumor infiltrate assessed for cDC1/cDC2 fractions. **(E)** Survival fractions of vehicle or αPD-L1 treated WT or *IRF8+32^-/-^* mice. Data are represented as mean +/- SEM of at least three independent experiments. **** p <0.0001. Differences in DC count analyzed with unpaired two-tailed T tests. Survival analyzed with log-rank test between individual groups.

We next tested the hypothesis that cDC1 are required to mount an immune response against GL261 brain tumors. We employed the *IRF8+32kb^-/-^* mouse model of selective cDC1 deficiency (*51*). This model possesses a 0.5kB deletion at the IRF8/Batf3 binding site within a super enhancer +32kb downstream from the *IRF8* transcription start site, which blocks cDC1 development (*22*). Compared to wild type mice, the immune infiltrate within GL261 tumors transplanted into *IRF8+32kb^-/-^* brains lacked XCR1^+^ cDC1 (Figure 1D). We next determined whether cDC1 were necessary for checkpoint blockade responsiveness. Although GL261 grows progressively in immune competent mice, treatment of tumor-bearing mice with anti-PD-1/PD-L1 improves survival (*52–55*). Consistent with prior work (*52–55*), we demonstrated that anti-PD-L1 treatment increased the median survival as well as the overall survival of wild type mice with GL261 brain tumors compared to vehicle-treated mice (Figure 1E). In contrast, cDC1-deficient *IRF8+32kb^-/-^* mice experienced no survival benefit when treated with anti-PD-L1 compared to vehicle-treated mice (Figure 1E). Control mice from both wild type and *IRF8+32kb^-/-^* mice experienced no difference in median or overall survival. These data show that DC (including the cDC1 subset) are recruited to the brain tumor microenvironment in numbers significantly higher than in the steady state brain parenchyma, and that cDC1 are required for response to anti-PD-L1 checkpoint blockade treatment of GL261-bearing mice.

### cDC1 prime CD8^+^ T cell responses against glioblastoma

Because cDC1 can present antigen to prime T cell responses, we determined the effects of cDC1 deficiency on T cell composition within the brain tumors of GL261-bearing mice. Compared to wild type mice, tumor infiltrating lymphocytes (TIL) of cDC1-deficient *IRF8+32kb^-/-^* mice harbored decreased numbers of all T cells, including both non-Treg CD4^+^ and CD8^+^ T cells, as a percentage of all tumor-infiltrating CD45^+^ cells (Figure 2A). In contrast, we observed similar numbers of CD4^+^ T regulatory cell percentages in GL261 TIL from both *IRF8+32kb^-/-^* and WT mice (Figure 2A). Moreover, a smaller proportion of CD8^+^ T cells directly isolated from GL261 tumors transplanted in *IRF8+32kb^-/-^* mice expressed the functional markers Granzyme B and PD- 1 compared to wild type mice (Figure 2B). These data show that GL261 tumors from *IRF8+32kb^-/-^* mice harbor fewer T cells, and a smaller proportion of the infiltrating CD8^+^ T cells exhibit an activated phenotype.

**Figure 2.**
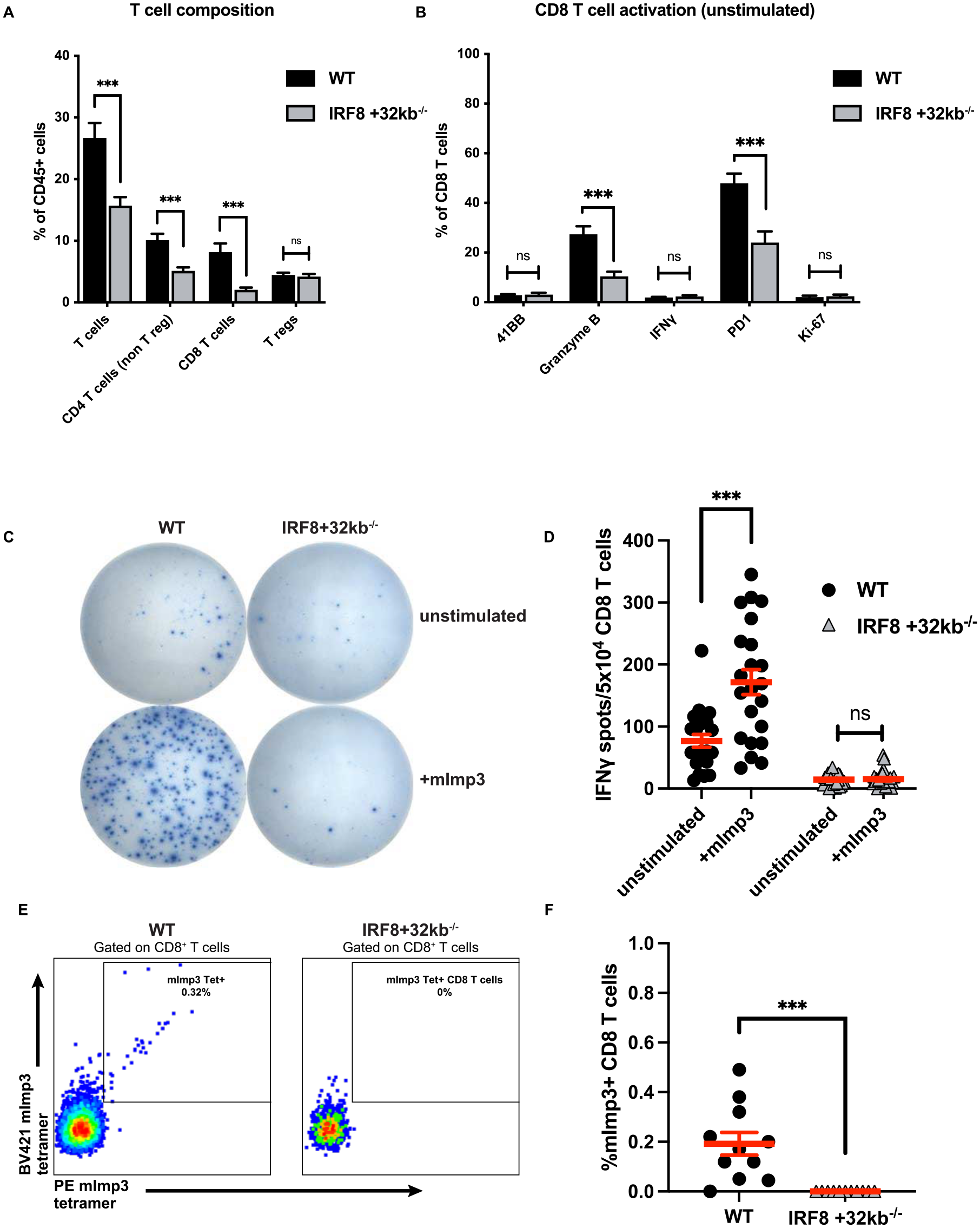
cDC1 prime effector and neoantigen-specific CD8 T cell responses against mouse glioma. **(A)** T cell composition of WT vs *IRF8+32kb^-/-^* GL261 brain tumors as assessed by flow cytometry. **(B)** CD8^+^ T cell activation marker expression of WT vs *IRF8+32kb^-/-^* GL261 brain tumors as assessed by flow cytometry. **(C)** mImp3 neoantigen specific CD8^+^ T cell response assessed by Interferon-γ release assay (ELISPOT) with graph **(D)**. **(E)** mImp3 neoantigen specific CD8^+^ T cell response assessed by tetramer staining and flow cytometry with graph **(F)**. Day 14 tumors for every panel in this figure. Data are represented as mean +/- SEM of at least three independent experiments. *** p <0.001. Differences in cell infiltrate, tetramer binding, and IFN-γ release analyzed with unpaired two-tailed T test.

Although studies have demonstrated that cDC1 cross-present antigen to prime CD8^+^ T cell responses in tumors outside the CNS (*36–43*), it remains unclear whether they perform a similar function during the immune response to brain tumors, particularly given the paucity of DC in steady-state brain parenchyma. Therefore, we tested the hypothesis that cDC1 are required to prime neoantigen-specific CD8^+^ T cells against GL261 brain tumors. Previously, we identified that mice harboring intracranial GL261 mount endogenous CD8^+^ T cell responses against the H- 2D^b^ restricted neoantigen, mutant Imp3 (mImp3) (*56*). In contrast to wild type mice, we identified that equal numbers of CD8^+^ T cells isolated from brain tumors of cDC1-deficient *IRF8+32kb^-/-^* mice bearing GL261 did not produce interferon gamma (IFN-γ) in response to stimulation with mImp3 peptide by ELISPOT (Figure 2C-D). Furthermore, *IRF8+32kb^-/-^* mice completely lacked mImp3-specific CD8^+^ T cells in the tumor infiltrate compared to wild type mice as assessed by staining with H-2D^b^-mImp3 tetramer (Figure 2E-F). These data show that cDC1 are required to prime effector CD8^+^ and neoantigen specific CD8^+^ T cell responses against GL261 brain tumors.

### Tumor antigen-containing cDC1 infiltrate the tumor and appear in cervical lymph nodes

One working model of cDC1 is that they phagocytize antigen in the periphery and home to secondary lymphoid tissues to prime naive T cells. Although the brain parenchyma does not harbor conventional secondary lymphoid tissue, compelling data have implicated the extracranial cervical lymph nodes as central to the priming of CNS antigen-specific T cells (*29, 32-34, 55, 57*). To determine if tumor antigen-containing cDC1 could be identified in these anatomic locations, we leveraged the characteristic that cDC1 can phagocytize target material, activate, and migrate to lymph nodes. We tested the hypothesis that fluorescent protein overexpressed by brain tumor cells could be phagocytized and retained by cDC1, and this “transfer of fluorescence” could be tracked by flow cytometry similar to previously employed approaches in pre-clinical models of melanoma (*39, 41, 58*).

We created two transgenic orthotopic glioma cell lines: GL261-zsGreen and CT2A-zsGreen. When transplanted into mice, the tumor-derived zsGreen protein can be phagocytosed by infiltrating innate immune cells and used as a detectable surrogate for tumor antigen uptake and trafficking (Figure 3A). Because CT2A-zsGreen tumors most consistently retained zsGreen expression at the time of harvest following orthotopic transplantation, we used this tumor for most tumor-antigen tracking experiments. To test the hypothesis that brain tumor-infiltrating cDC1 contain tumor antigen, we injected CT2A-zsGreen tumors intracranially and monitored for the presence of zsGreen within cDC1, as well as other immune cells, by flow cytometry. In CT2A- zsGreen tumors, we identified zsGreen present not only within cDC1 but also cDC2, monocyte derived DC (MoDC), and plasmacytoid DC (pDC) infiltrating the tumor (Figure 3B), suggesting that these populations phagocytose tumor-derived antigens *in vivo*. We did not identify zsGreen^+^ APC within non-fluorescent CT2A tumors. We then examined the ipsilateral cervical lymph nodes (cLN) to determine whether zsGreen^+^ APC were present extracranially. We identified zsGreen^+^ cDC1 (both migratory and resident subsets) in both the deep and the superficial cLNs (Figure 3C- D). A larger fraction of migratory cDC1 (further defined by CD103^+^CD8α^-^) were zsGreen^+^compared to the resident cDC1 subset (further defined by CD8α^+^CD103^-^). We also identified other zsGreen^+^ APC populations, including cDC2, MoDC and pDC in the superficial cLNs, and MoDC in the deep cLNs. Although GL261-zsGreen tumors less reliably retained zsGreen expression *in vivo*, zsGreen was still detectable within a considerable proportion of the tumor infiltrating DC (Figure S1A), as well as within APC subsets isolated from the deep and superficial cervical lymph nodes. We detected a significant proportion zsGreen^+^ cDC2 in the deep cervical lymph node, as well as cDC2, MoDC and pDC in the superficial cervical lymph node (Figure S1B-C). This extended our findings with CT2A zsGreen brain tumors. These data show that cDC1 as well as other innate immune cell populations harbor tumor-derived antigen both within the tumor microenvironment and also in the cervical lymphoid tissue across different brain tumor preclinical models.

**Figure 3.**
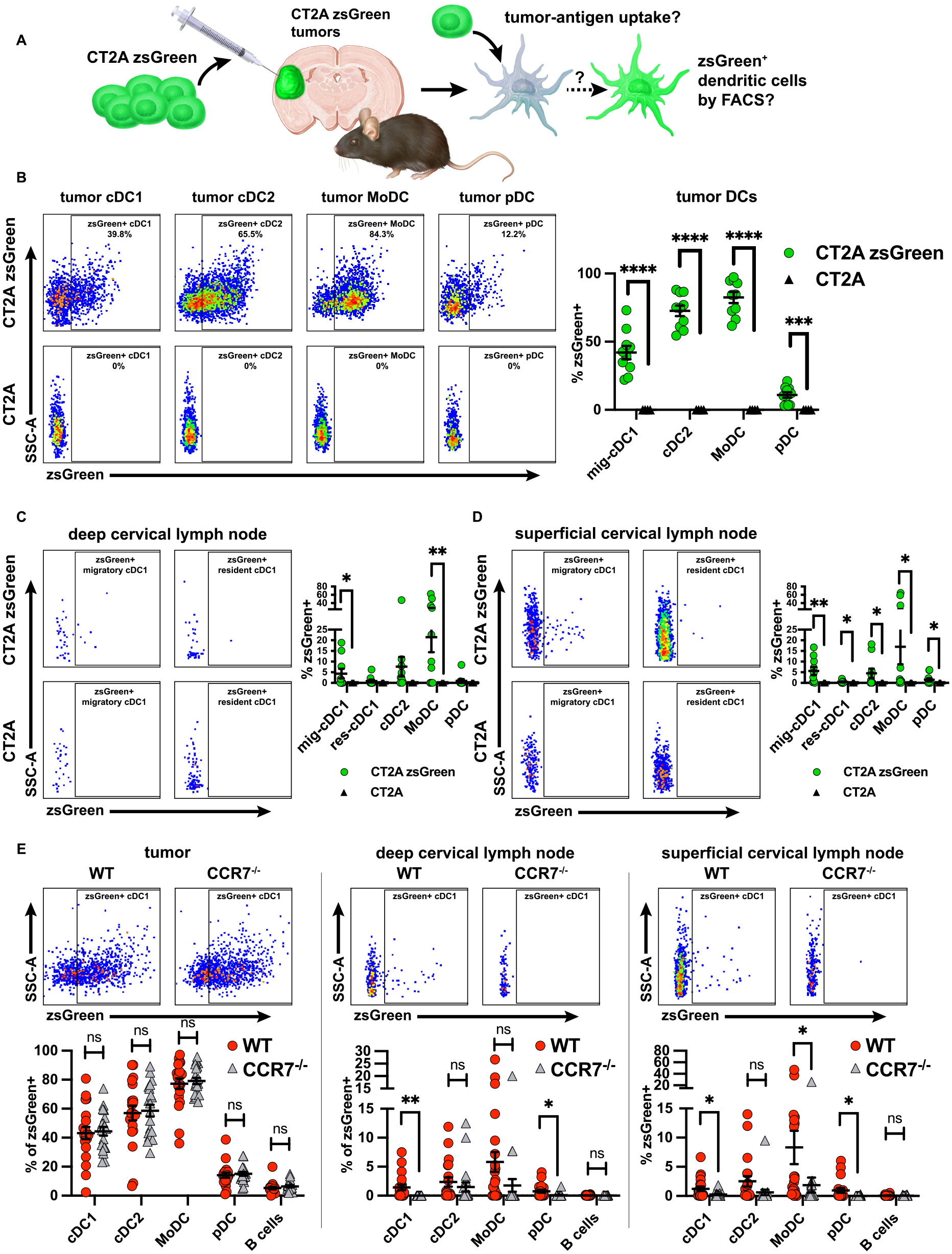
cDC1 derived from cervical lymph nodes harbor tumor-derived antigen, and do so in a CCR7-mediated manner. **(A)** schematic describing principles and process. **(B)** zsGreen phagocytosis by DC infiltrating the CT2A-zsGreen tumor. zsGreen expression by **(C)** deep cervical lymph node and **(D)** superficial cervical lymph node DC. **(E)** zsGreen phagocytosis by DC infiltrating the CT2A zsGreen tumor of WT vs CCR7^-/-^ mice. zsGreen expression by **(F)** deep cervical lymph node and **(G)** superficial cervical lymph node DC of WT vs CCR7^-/-^ mice. Day 14 tumors for B-D, Day 12 tumors for (E) Data are represented as mean +/- SEM of at least three independent experiments. * p<0.05, **p<0.01, ***p<0.001, ****p<0.0001. Differences in %zsGreen^+^ analyzed with unpaired one-tailed T test with Welch’s correction. Grubb’s outlier test in (E) used to test for and exclude an outlier from both the WT and the *CCR7*^-/-^ genotypes.

Having observed zsGreen within cervical lymph node cDC1, we considered several scenarios by which tumor-derived antigen is trafficked from the CNS to cervical lymph nodes. Specifically, brain tumor-associated antigens could be actively trafficked by cellular transport within APC, or passively transferred from the brain to the cLN via acellular mechanisms. Regarding the former, migratory cDC1 home to draining lymph nodes via chemokine receptor 7 (CCR7)-mediated chemotaxis (*59*). Therefore, we tested the hypothesis that CCR7 expression was required for zsGreen^+^ cDC1 to appear in the cLN. Because *CCR7^-/-^* mice lack migratory DC in lymph nodes, we did not distinguish between migratory and resident cDC1 in our analysis, and instead included all cDC1. We transplanted CT2A-zsGreen into the brains of both WT and *CCR7^-/-^* mice and monitored the zsGreen signal in immune cells isolated from brain tumors as well as from cLN. We observed equivalent zsGreen uptake by tumor infiltrating cDC within both WT and *CCR7^-/-^* mice, suggesting that cDC originating from mice of both genotypes were able to infiltrate brain tumors and phagocytose tumor antigen (Figure 3E, left). However, we observed a significantly attenuated zsGreen signal within the cDC1 isolated from the deep and superficial cervical lymph nodes of *CCR7^-/-^* mice compared to WT mice (Figure 3E, middle and right). Small but equal fractions of B cells were positive for zsGreen in cervical lymph nodes of wild type and *CCR7^-/-^* mice (Figure 3E, middle and right), which may suggest that passive CNS lymphatic drainage is possible in both genotypes since B cells are not known to migrate from the periphery to lymph nodes, and instead enter the lymph node through high endothelial venules (*60–62*). The zsGreen signal was incompletely extinguished in DC within *CCR7^-/-^* lymph nodes, which suggests that passive drainage or a CCR7-independent process could be potential mechanisms to traffic a small fraction of tumor antigen to draining lymph nodes. The diminished lymph node zsGreen signal in *CCR7^-/-^* mice was not restricted to the cDC1 subset, and included the pDC in both superficial and deep cLNs, and MoDCs in superficial cLNs (Figure 3E, middle and right). Together with our previous observation that cDC1 are required to generate brain tumor-specific T cell responses, these results suggest that cDC1 perform this function, at least in part, by phagocytizing tumor-associated material within the brain and trafficking it to the lymph nodes in a CCR7/cell migration-dependent manner.

### Dura associated cDC1 undergo dynamic changes in response to GBM

Having demonstrated that cDC1 harbor tumor-derived antigen within brain tumors as well as in the draining cervical lymph nodes, we next characterized cDC1 within cranial dura, the outer fibrous layer which surrounds the brain. The dura has been increasingly recognized as an immunologically dynamic tissue site. While the steady state brain parenchyma is devoid of cDC1, dura harbors a diverse composition of leukocytes including conventional DC, plasmacytoid DC, macrophages, T cells, and B cells, among other cell types (*22, 63*). The dura also contains lymphatic vessels that traverse the venous sinuses and drain to the cervical lymph nodes (*28, 29, 31, 55*). Prior studies have demonstrated the presence of Flt3L responsive DC in the dura (*22, 23*). Given the growing recognition of the importance of the dura microenvironment during immune responses, we addressed both the location of the specific cDC1 subset as well as the antigen presentation capacity of dura-associated cDC during brain tumor growth.

We investigated the location of dura-associated cDC1 in *SNX22^GFP/+^* mice, in which cDC1 specifically are labeled with GFP, by 2-photon microscopy. In control sham-treated and GL261- OFP bearing *SNX22^GFP/+^* mice, the dura harbored extravascular cDC1 (Figure 4A). We hypothesized that the number of cDC1 infiltrating the dura would increase in response to the presence of a tumor in the brain parenchyma. Compared to sham-injected mice, dura from intracranial GL261-bearing mice contained increased numbers of cDC1, cDC2, MoDC and pDC by flow cytometry (Figure 4B).

**Figure 4.**
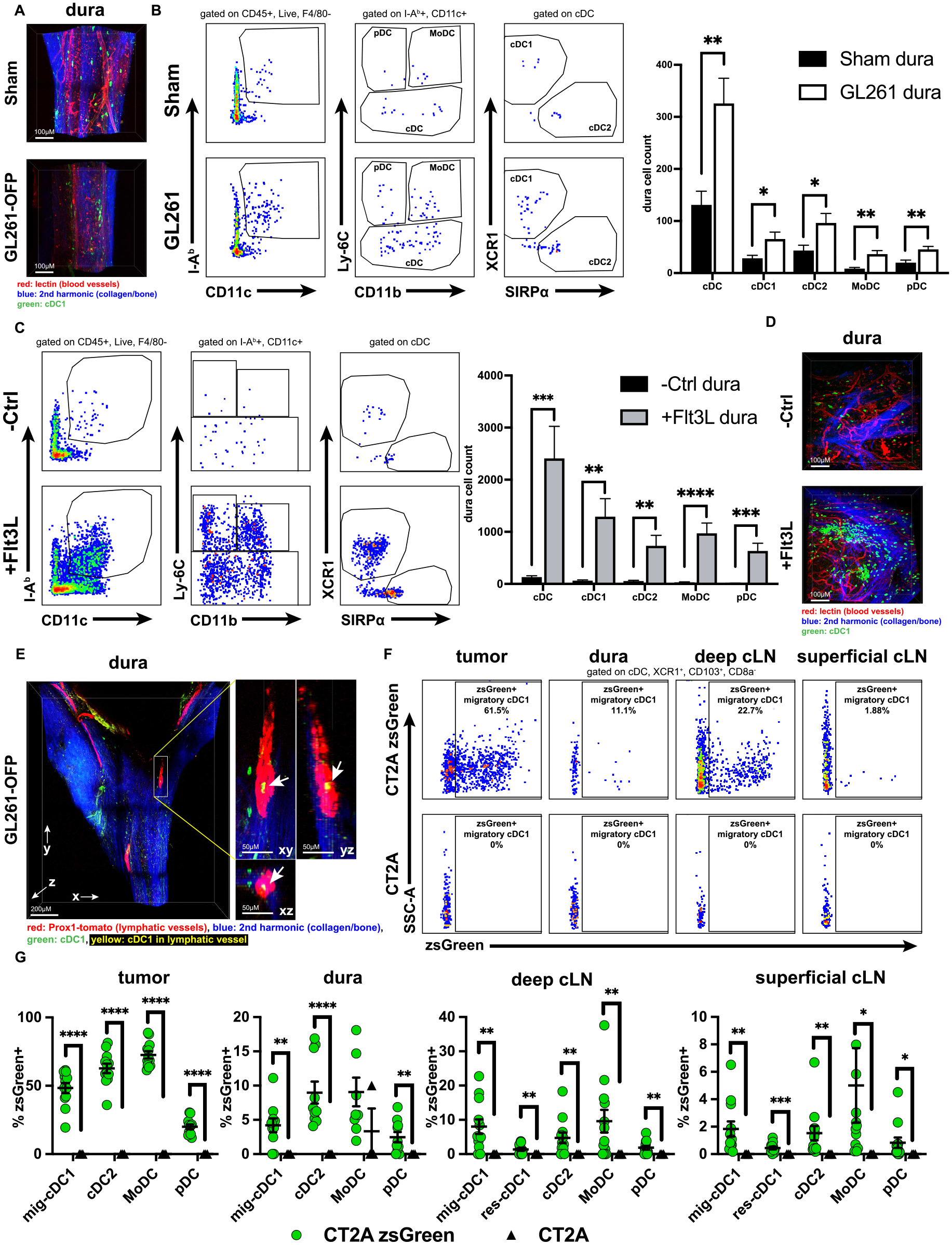
Dendritic cells harbored by dura and dura lymphatics and undergo dynamic changes in response to intracranial tumors. **(A)** 2-photon microscopy of *SNX22^GFP/+^* mouse dura along the superior sagittal sinus depicting sham vs. GL261 OFP intracranially injected mice. Dura DC assessed and quantified by flow cytometry in **(B)** sham injected vs. GL261 injected brains at d14 post tumor injection, and **(C)** -Ctrl vs Flt3L treated mice. **(D)** 2-photon microscopy of *SNX22^GFP/+^* mouse dura depicting -Ctrl vs Flt3L treated mice. **(E)** Dura from intracranial GL261- OFP bearing tamoxifen treated *SNX22^GFP/+^/Prox1-Cre-tdTomato^+/-^* mice, with GFP^+^ cDC1 and tdTomato^+^ lymphatic vessels near bregma—cDC1 visualized in lymphatic vessel. **(F)** Dura associated zsGreen^+^ migratory cDC1 assessed by flow cytometry, and **(G)** quantified across all DC subsets. Dura samples with CD45^-^, zsGreen^+^ cells were assumed to be contaminated by tumor infiltrate, and were excluded from analysis. Data are represented as mean +/- SEM of at least three independent experiments. * p<0.05, **p<0.01, ***p<0.001, ****p<0.0001. Differences in DC count analyzed with unpaired two-tailed T tests. Differences in %zsGreen^+^ analyzed with unpaired one-tailed T test with Welch’s correction.

We examined whether the growth factor responsible for conventional DC development, Flt3L, could also be used to also drive the expansion of the cDC1 subset specifically within the dura. Previous work showed that dura harbors CD11c^+^, I-A^b+^ cells whose expansion could be driven by systemic Flt3L administration (*23*), and a recent study identified that systemic Flt3L administration could drive expansion of cDC1 and cDC2 when the whole brain and surrounding meninges were examined together (*22*). We extended these data and focused specifically on the effect of Flt3L on dura cDC1. When mice were administered systemic Flt3L, we observed significant expansion of DC subsets in the dura, including the cDC1 subset, by flow cytometry in wild type mice (Figure 4C) and 2-photon microscopy in the *SNX22^GFP/+^* mouse (Figure 4D, Figure S2A). Flt3L induced a greater magnitude of expansion of dura DC than intracranial-GL261. These data show that dura-resident cDC1 can undergo expansion driven both by the presence of tumor located within the brain parenchyma as well as by systemically-administered Flt3L.

To investigate the spatial relationship between cDC1 and dura lymphatic vessels, we crossed the *SNX22^GFP/GFP^* cDC1 reporter mouse with the *Prox1-Cre-tdTomato^+/+^* reporter mouse, which expresses tdTomato specifically in lymphatic vessels (*64*). Tamoxifen-treated F1 mice have GFP^+^ cDC1 and tdTomato^+^ lymphatic vessels. Using 2-photon microscopy to characterize the dura in tamoxifen-treated F1 mice, we observed that GFP^+^ cDC1 could be detected within the dura lymphatic vessels themselves when mice were orthotopically transplanted with intracranial GL261-OFP (Figure 4E, Figure S2B) or were sham-injected (Figure S2C). These data show that cDC1 can be identified within the lumen of dura-lymphatic vessels from both the tumor-bearing and steady-state brain conditions.

Having identified cDC1 within the dura and, in some instances, within dura lymphatic vessels, we tested the hypothesis that dura cDC1 could acquire tumor-derived antigen from intraparenchymal brain tumors. Because CT2A-zsGreen retained the highest levels of zsGreen expression at the time of harvest compared to GL261-zsGreen, we used CT2A-zsGreen to monitor for tumor-antigen presence in dura-associated cDC1. We orthotopically transplanted CT2A-zsGreen tumors into the brain parenchyma of mice and monitored the zsGreen signal within immune cells on day 7 post-injection within tumors, dura, and cervical lymph nodes by flow cytometry. To avoid contamination of the dura with underlying parenchymal zsGreen^+^ tumor and tumor infiltrating immune cells, we (a) selected an early time point following tumor injection to ensure that tumors were small and did not abut the dura and (b) resected the dura surrounding the prior injection site to avoid potential contamination of the dura samples by adjacent tumor and immune cells. We observed zsGreen^+^ migratory cDC1 in the tumor itself, the dura, the superficial cervical lymph nodes, and the deep cervical lymph nodes (Figure 4F). Importantly, we did not observe a CD45^-^, zsGreen^+^ population in the dura, which indicates that our dura samples were not contaminated by tumor or tumor infiltrate. We also observed zsGreen in cDC2, pDC, and monocyte-derived DC in the tumor, dura, and cervical lymph nodes (Figure 4G). Together, these data show that dura-associated Flt3L-responsive cDC1 can localize to dura lymphatic vessels and that dura-associated cDC1, along with other dura-associated APC subsets contain tumor antigen from intraparenchymal tumors.

### CD8^+^ T cell priming occurs in cervical lymph nodes and requires cDC1 and CCR7 for normal kinetics

Having established that tumor-antigen containing DC can be found in the dura as well as the superficial and the deep cervical lymph nodes, we examined where T cell priming takes place during the immune response to CNS tumors. The location of T cell priming during immune responses against CNS tumors remains incompletely characterized. Several observations support a role for the cervical lymph nodes in CNS pathology: (1) ligation of deep cervical lymph nodes attenuated the effects of VEGF-C-mediated elimination of brain tumors (*55*); (2) lymphadenectomy of cervical lymph nodes ameliorated the disease burden in rodent models of EAE (*65–67*); and (3) ablation of meningeal lymphatics decreased the antigen specific T-cell-CD11c+ interactions in deep cervical lymph nodes of mice with EAE (*68*). We extended these studies to address where T cell priming might take place in the immune response against CNS tumors.

Because we detected tumor-antigen containing APCs in several anatomic sites, we considered the possibilities that CD8^+^ T cell priming could occur within the tumor itself, the dura, the cervical lymph nodes, or the spleen. To this end, we tracked the cell division *in vivo* of OT-I T cells adoptively transferred into mice bearing intracranial GL261 cells transduced with full length ovalbumin (GL261-OVA). OT-I T cells recognize the H-2K^b^ restricted ova antigen SIINFEKL (*69*) and proliferate upon being primed. We transplanted GL261-OVA into the brains of CD45.2^+^ C57BL/6 hosts. Four days post-transplant, we adoptively transferred CFSE-labeled, CD45.1^+^CD45.2^+^ congenically marked OT-I CD8^+^ T cells. We harvested the region of the cerebral hemisphere which contained the transplanted tumor cells, the dura, the ipsilateral superficial and deep cervical lymph nodes, the spleen, and the non-CNS draining contralateral inguinal lymph node, and monitored for CFSE dilution of OT-I CD8^+^ T cells within these disparate tissues (Figure 5A-C). We distinguished between CFSE-low, CFSE-mid, and CFSE-high OT-I CD8^+^ T cells which we envisaged to reflect terminal effector cell function, division during initial priming, or no division due to the absence of priming, respectively. We harvested tissues at days 3 and 6 post adoptive transfer to capture early and late snapshots of the evolution of clonal expansion within each of the tissues.

**Figure 5.**
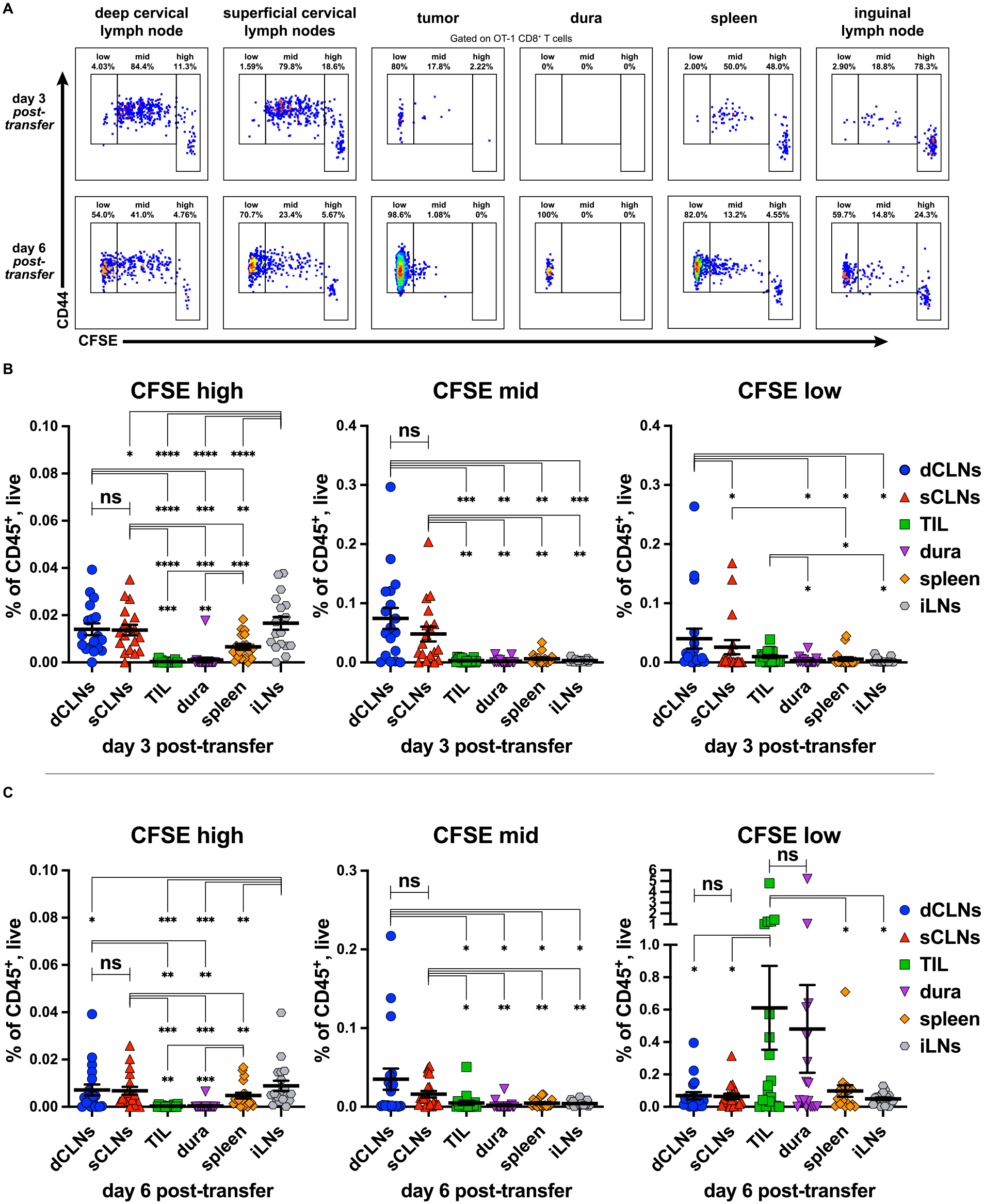
Clonal Expansion of OT-I CD8^+^ T cells occurs primarily in extracranial CNS-draining cervical lymph nodes. **(A)** CD44 expression and CFSE dilution of OT-I CD8^+^ T cells assessed by flow cytometry at day 3 and day 6 post adoptive transfer of ipsilateral superficial cervical lymph nodes, ipsilateral deep cervical lymph nodes, cerebral hemisphere region encompassing tumor, dura (with tumor abutting region resected), spleen, and non-draining contralateral inguinal lymph node. Quantitation of CFSE high, mid, and low OT-I CD8^+^ T cells at **(B)** day 3 and **(C)** day 6 post-transfer. Data are represented as mean +/- SEM of at least three independent experiments. * p<0.05, **p<0.01, ***p<0.001, ****p<0.0001. Differences between organs of OT-I CFSE low/mid/high % of CD45^+^/Live compared with paired two-tailed T tests.

At 3 days post-adoptive transfer, we observed the greatest numbers of CFSE-high, undivided OT-I cells as a fraction of the CD45^+^ compartment in the lymph nodes, regardless of location (Figure 5B, left). The tissue with the next largest fraction of undivided OT-I cells was the spleen (Figure 5B, left). The tumor and the dura had almost no CFSE-high, undivided OT-I T cells (Figure 5B, left), commensurate with our understanding that naive T cells circulate between lymphoid organs before activating and dividing. This same pattern held through day 6 post-adoptive transfer, although there were fewer undivided OT-I cells as a fraction of CD45^+^ cells (Figure 5C, left). At day 3 post-adoptive transfer the CNS-draining superficial and deep cervical lymph nodes harbored significantly more CFSE-mid initially divided OT-I T cells as a fraction of the CD45^+^ compartment than other tissues (Figure 5B, middle). This is consistent with our understanding that clonal expansion first occurs in proximal draining lymph nodes. This same pattern held through day 6 post-transfer, although there were fewer CFSE-mid OT-I cells as a fraction of CD45^+^ cells (Figure 5C, middle). At day 3 post-adoptive transfer, CFSE-low terminally divided OT-I T cells occupied the greatest fraction of CD45^+^ cells in the deep cervical lymph nodes but were also observed in the superficial lymph nodes and TIL, albeit at lower numbers (Figure 5B, right). However, by day 6 post-transfer, this pattern had shifted. The TIL harbored the greatest fraction of terminally divided OT-I T cells as a fraction of CD45^+^ cells compared to all other sites except the dura (Figure 5C, right). This underscores our understanding that the tumor is both the source of antigen and effector site for OT-I T cells. We observed similar patterns of expansion of OT-I cells in CT2A-OVA tumors, although the expansion was weaker and somewhat delayed when compared with GL261- OVA experiments (Figure S3A), which may be due to the immune-suppressive effects of CT2A. Together, these data support a model in which CD8^+^ T cells are primed against brain tumors in the CNS-draining cervical lymph nodes, and subsequently home to the CNS to mount an anti-tumor immunity there, and that, while the tumor itself is a primary site of effector function, effector T cells can also be observed in the dura.

Given our observations that cDC1 are required to prime neoantigen specific T cell responses against CNS tumors, and that CCR7 is required to traffic antigen from brain tumor to draining cervical lymph nodes, we next investigated their role in clonal expansion of adoptively transferred OT-I T cells in mice with GL261-OVA brain tumors. Compared to WT mice, *CCR7^-/-^* and *IRF8+32kb^-/-^* mice had decreased clonal expansion of OT-I T cells in the cervical lymph nodes and spleen at 3 days post adoptive transfer (Figure S4A, 1st, 2^nd^, and 5^th^ graph from left). Notably, in contrast to day 3, at day 6 we observed minimal differences in clonal expansion between WT, CCR7^-/-^, and IRF8+32kb^-/-^ mice (Figure S4B). These data collectively demonstrate that CD8^+^ T cell priming and clonal expansion against brain tumor-specific antigens occurs in the cervical lymph nodes, and that absence of intact cell migration to lymph nodes and cDC1 deficiency lead to delayed, but not completely absent clonal expansion in this setting.

### The CD141^+^ human equivalent of the cDC1 is detectable in human in the dura and brain tumors

Having identified cDC1 and additional cDC subsets within murine brain tumors and matched dura, we investigated whether human tumors and dura also harbored infiltrating DC populations. While human DC subsets and functions are less well characterized than in mice, the human cDC1 parallel is the CD141-expressing cDC. In both mice and humans, experiments have demonstrated that these parallel cDC1 subsets can produce IL-12 (*70*), cross present exogenous proteins derived from internalized dead cells to CD8^+^ T cells (*70*), and express high levels of IRF8 (*71*), a critical regulatory factor required for cDC1 development in mice (*51*) and in humans (*72*).

We explored the immune cell populations within the tumor and matched dura of patients undergoing craniotomies for tumor resection in which, in addition to the tumor, there was a clinical indication to also resect adjacent normal, non-tumor involved dura (Figure 6A). We disaggregated dura and matched tumor specimens from five meningioma patients and one GBM patient and performed flow cytometry. In both GBM and meningiomas, we detected the presence of multiple human DC equivalents vested within tumor and matched dura specimens, including cDC1- equivalent CD141^+^ DC, cDC1-equivalent CD1c^+^ DC, as well as CD14^+^ monocytes, and CD16^+^ monocytes (Figure 6B-D). We identified cDC1, cDC2, CD14^+^ monocytes, CD16^+^ monocytes, CD4^+^ T cells, and CD8^+^ T cells in most of the thirteen total GBM tumors that we analyzed (Figure 6C). Moreover, between GBM and meningiomas, the matched dura samples harbored similar fractions (as a percentage of CD45^+^ cells) of each of the above-mentioned cell types, although our analysis was limited to a single GBM sample in which dura was also resected (Figure 6F). These findings collectively show that human conventional DC subsets as well as monocytes are abundantly detectable in dura and tumors and across multiple brain tumor types, which suggests they could play a role in human CNS anti-tumor immunity.

**Figure 6.**
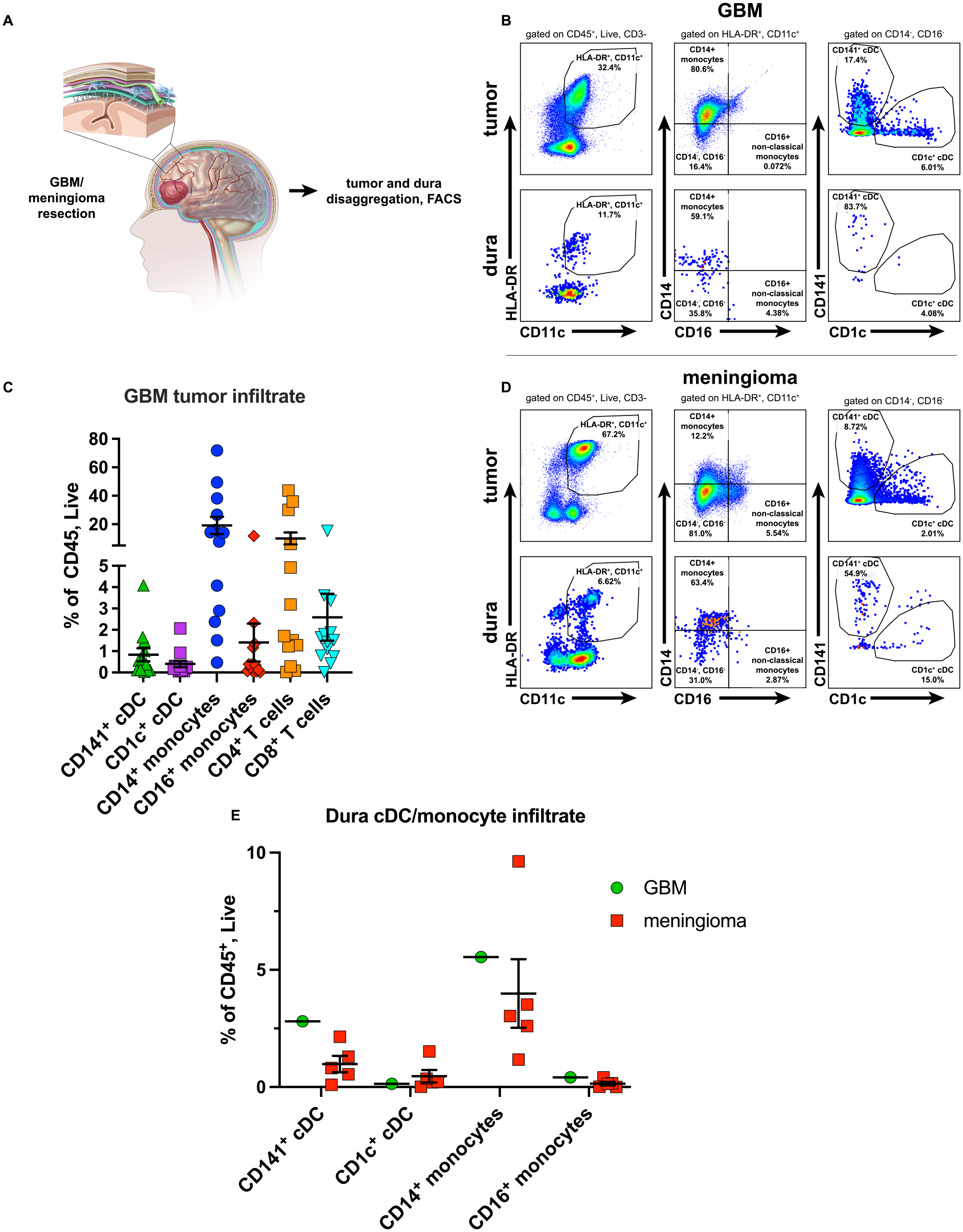
Dendritic cells infiltrate human dura and tumors. **(A)** Representative sketch of resection of brain with meningioma/GBM and associated dura (also resected when clinically indicated). Dura and tumor infiltrating CD141^+^ cDC and CD1c^+^ cDC assessed by flow cytometry of **(B)** a GBM with **(C)** associated immune cell infiltrate composition as well as **(D)** a meningioma. **(E)** Dendritic cell and monocyte composition between a GBM-associated dura sample and meningioma associated dura samples.

### The CD141^+^ cDC and other APC phagocytose a tumor-specific marker in GBM

We next applied the “transfer of fluorescence” methodology employed previously to clinical studies of brain tumors from patients in order to characterize the immune cells that phagocytize GBM-derived material. Similar to the preclinical studies in which we tracked tumor-derived zsGreen, this approach relies on the premise that fluorescent material derived from the tumor could potentially be transferred to, and detectable within, DC subsets and other phagocytic cells infiltrating the tumor. During GBM resection, the FDA-approved compound 5-aminolevulinic acid (5-ALA) can be used in “fluorescence-guided surgery” to fluorescently label the tumor in order to distinguish it from normal brain (*44–47*). After patients ingest 5-ALA preoperatively, the compound crosses the blood-brain barrier and is metabolized by target GBM cells to protoporphyrin IX (PPIX) (*48*). The accumulation and retention of PPIX is highly specific to GBM cells (*48, 49*). During GBM resection, illumination of the tumor with blue light from an operating microscope leads to fluorescent excitation of the GBM-associated fluorophore PPIX, which emits light within pink wavelengths. In contrast to non-fluorescent normal brain tissue, tumor tissue fluoresces pink under the microscope (Figure 7A). We tested the hypothesis that a subset of phagocytic CD45^+^ cells, including DC, could acquire and retain tumor-derived PPIX in patients undergoing 5-ALA-mediated fluorescence guided surgery for GBM (Figure 7B).

**Figure 7.**
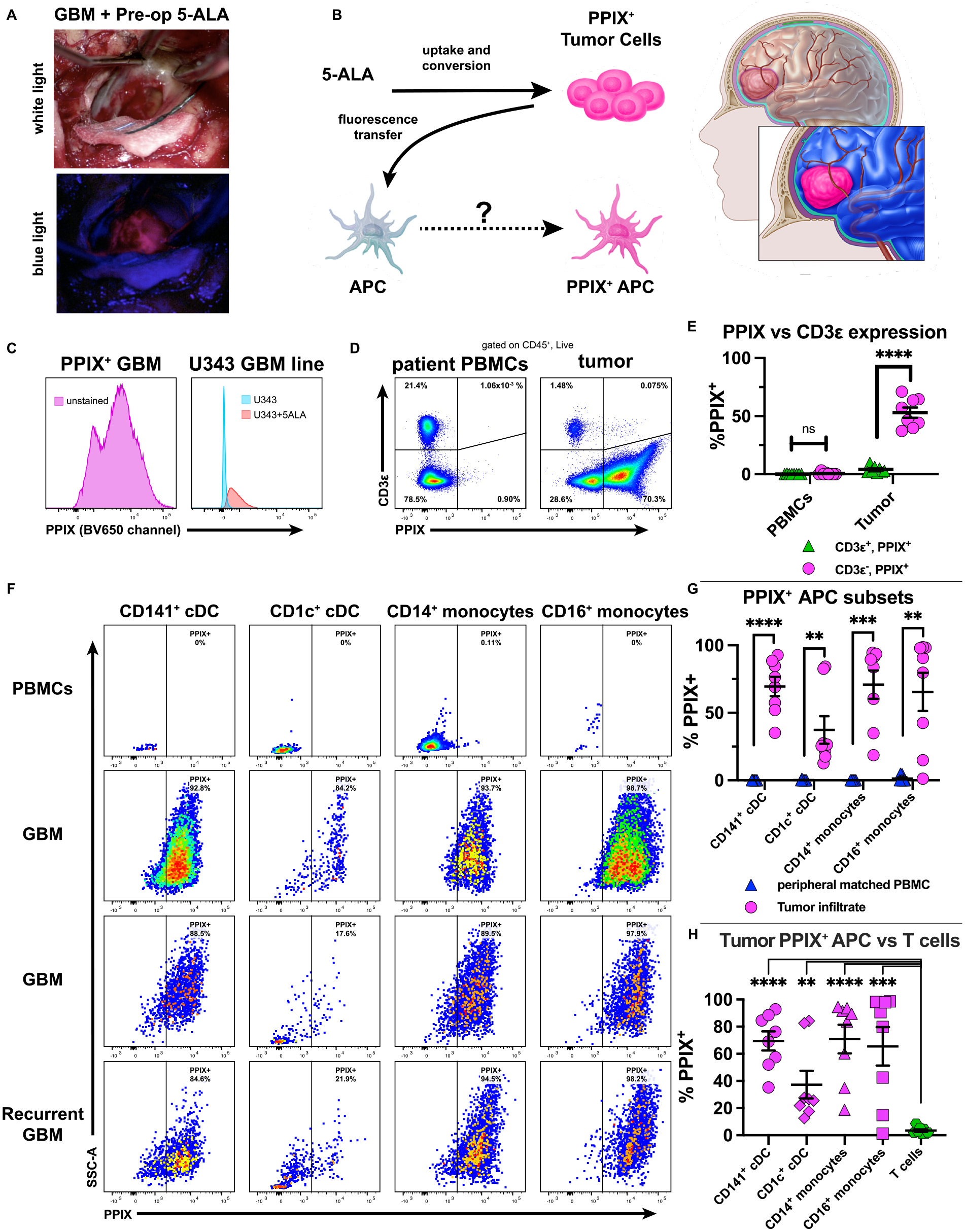
APC infiltrating human GBM uptake a tumor-specific reporter PPIX. **(A)** GBM patient with pre-administered 5-ALA: tumor visualized under white (top) or blue (bottom) light. **(B)** Schematic describing principles and process. **(C)** PPIX expression in either a bulk non-antibody-stained tumor from a 5-ALA resected GBM (left) or in U343 cells treated with 5-ALA (right). **(D)** CD3ε vs PPIX expression of CD45^+^/Live cells derived from patient intraoperatively derived PBMC (left) or resected GBM tumor (right) with associated graph **(E)**. **(F)** PPIX^+^ APC subsets across 3 GBM tumors (two primary, one recurrent) compared to peripheral intraoperatively derived patient PBMC with associated graph **(G)**. **(H)** Graph of tumor infiltrating PPIX^+^ APC vs. T cells. Data representative of eight patients (six primary, two recurrent) in which GBM and matched intraoperative PBMC were taken. **p<0.01, ***p<0.001, ****p<0.0001. Unpaired two-tailed T tests for (E) and (H). Paired two-tailed T tests for (G).

To test whether we could detect PPIX^+^ cells by flow cytometry, we characterized samples both form 5-ALA-treated patients as well as *in vitro* passaged cell lines exposed to 5-ALA. We disaggregated a GBM specimen obtained from a 5-ALA-mediated surgical resection and performed flow cytometry on the single cell suspension. We determined that the PPIX signal was brightest in the BV650 channel (Figure S5). We also observed clearly defined PPIX-low and PPIX- high cell populations, which suggests a range of PPIX uptake and retention amounts among tumor cells (Figure 7C, left). We thus used the BV650 channel to detect PPIX for further experiments. We also treated the U343 glioblastoma cell line with 5ALA and performed flow cytometry. Compared to untreated U343 cells, 5-ALA treated U343 cells also fluoresced brightly in the BV650 channel (Figure 7C, right).

We next analyzed 5-ALA treated GBM patient samples to determine if transfer of PPIX fluorescence from tumors to infiltrating APC occurred in patients. Specifically, we addressed two central questions: (1) would transfer of PPIX fluorescence from tumor to immune cells occur only in the tumor microenvironment and not in the periphery? and (2) within the tumor mass, would only phagocytic immune cells acquire and retain tumor-derived PPIX? To answer these questions, we studied the matched peripheral blood and tumor obtained from patients with GBM undergoing 5-ALA fluorescence guided surgery. We characterized both disaggregated tumor samples and patient-matched intraoperatively harvested peripheral blood mononuclear cells (PBMC) by flow cytometry. In PBMC, neither the CD3ε^+^ fraction, nor the CD3ε^-^ fraction had a detectable PPIX signal (Figure 7D, upper graph). In contrast, whereas the CD3ε^+^ fraction of the tumor infiltrate was PPIX-negative, a majority of cells in the CD3ε^-^ fraction were PPIX-positive (Figure 7D-E). These data show that presence of PPIX within an immune cell depends on both cell identity and localization within the tumor.

We next investigated whether tumor infiltrating DC harbored detectable PPIX. We disaggregated GBM tumors that were resected using 5-ALA. Compared to intraoperatively harvested PBMCs as a control, we observed PPIX within different APC subsets infiltrating the tumor, including CD141^+^ cDC, CD1c^+^ cDC, CD14^+^ classical monocytes, and CD16^+^ non-classical monocytes across eight GBM samples: six primary and two recurrent GBMs (Figure 7F-G). Moreover, the PPIX-signal localized to APC and not T cells infiltrating the tumor (Figure 7H). Thus, multiple tumor-infiltrating APC subsets acquire and retain a GBM-derived tracer. Together, these data show that PPIX transfer of fluorescence from tumor to infiltrating CD45^+^ cells depended on both location and cell identity. Phagocytic immune cells, and not T cells, acquired and retained PPIX. Moreover, only immune cells within the tumor microenvironment and not in the periphery became PPIX^+^, which underscores that the presence of CD45^+^PPIX^+^ cells was a function of both cell identity and location within the tumor.

## DISCUSSION

Here, we describe the functional importance of cDC1 in the immune response to GBM. We showed that cDC1 infiltrate GL261 and CT2A tumors and that they are critically required for the development of functional neoantigen-specific CD8 T cell responses to GBM as well as for responsiveness to checkpoint blockade in a model sensitive to this therapy. Moreover, using a traceable fluorescent tumor antigen model, we observed that multiple DC subsets, including cDC1, acquire tumor-derived material which is trafficked to both the superficial and deep cervical lymph node chains in a manner that requires CCR7 expression. We also identified cDC1 within the dura and observed that a subset of these cells are located within dura lymphatic vessels. Interestingly, we also detected tumor antigen within DC harbored by the dura, which could be a midpoint along the migration path of cDC1 transiting between the brain-tumor parenchyma and cervical lymph nodes. Based on the dynamics of expansion of T cells specific for an ectopic GBM antigen, lymph nodes were the predominant location of priming an immune response to GBM, and this process required intact cell migration and cDC1 for normal kinetics. Finally, we extended our work in mice to human tumors and dura. Both tumors and dura harbored the mouse cDC1-equivalent CD141^+^ DC, among other subsets. Moreover, in tumors from surgical resections in which 5-ALA was used to illuminate the tumor, APC that had infiltrated the tumor, but not the same subset from the periphery, harbored the fluorescent protoporphyrin IX tumor-antigen surrogate.

Previous reports have identified the critical role of cDC1 in anti-tumor immunity for a range of tumor types (*36–43*). Our findings extend the importance of the cDC1 to the immune response to CNS tumors. Recent work has indicated the importance of cDC1 in certain contexts within CNS anti-tumor immunity: investigators identified that the more immunogenic FGL2^-/-^ orthotopic brain tumors required Batf3-dependent cDC1 in order to be eliminated by the immune system (*73, 74*). The consequences of cDC1 deletion in our preclinical model extend their work and suggest that cDC1 are a required APC within the context of CNS anti-tumor immunity broadly, particularly to generate CD8^+^ Τ cell responses against the tumor. Further work will be necessary to understand the requirements for antigen presentation to CD4^+^ T cells in this setting. Nevertheless, despite the immunologically specialized nature of the CNS—absence of steady state parenchymal dendritic cells, lack of lymph nodes in the parenchyma, the presence of cerebrospinal fluid, and the blood brain barrier, among other features—the dependence on the cDC1 subset for immune responses to cancer appears to be shared between the CNS and other extracranial tissue sites.

The nature of antigen presentation in the CNS has been a topic of substantial investigation in several distinct disease models. Unlike other anatomic tissue sites, the only leukocytes present in any abundance in the brain parenchyma are the microglia (*22, 75*). In addition, border associated macrophages exist along the basement membranes of blood vessels (*22*). Although these innate cell types harbor the capacity to present antigen, they lack the capacity to migrate outside the CNS. In contrast, cDC possesses the canonical professional APC abilities to phagocytose antigenic material and migrate to secondary lymphoid tissue to prime naive T cells. In the CNS, cDC are relatively restricted to specialized anatomic locations including the choroid plexus and meninges (*22, 23*). Nevertheless, cDC can infiltrate the brain parenchyma in settings of inflammation (*24, 33, 34, 57, 63, 76, 77*). Previous studies from the Fabry group showed that intracerebrally injected monocyte-derived DC could migrate to the cervical lymph nodes (*32*), and EAE disease severity was attenuated when cDC subsets were depleted using the ZBTB46-DTR mouse (*76*). Other studies have shown that partial depletion of DC led to attenuated responses against GBM when anti PD-1 and/or PolyIC was used to stimulate the immune system to eliminate orthotopic models of GBM (*78*). Nevertheless, we have had a persistently limited understanding of how endogenous cDC1, rather than exogenously introduced dendritic cells, contribute to the immune response to brain tumors. Our work suggests that the cDC1 subset functions, at least in part, as a key professional APC in the immune response to brain tumors.

The relationship between CNS pathophysiology and the cervical lymph nodes has been supported by a growing body of work in different disease models. Extirpation of the cervical lymph nodes lessens EAE severity (*65–67*), and studies by both the Kipnis and Alitalo groups demonstrated the role of the cervical lymph nodes in lymphatic drainage of the CNS (*28, 29*). Recently, Iwasaki and colleagues demonstrated the critical involvement of the deep cervical lymph nodes in the VEGF- C mediated immune response to brain tumors (*55*). Given the importance of cervical lymph nodes in lymphatic drainage of the CNS, we explored how dendritic cells themselves might play a role in lymphatic drainage of the CNS, with particular attention paid to the endogenous mechanism of antigen trafficking. Similar fluorescence transfer approaches to the ones we employed have been used to show that DC in draining lymph nodes contain tumor-associated fluorescent antigen (*39–41, 58*). Prior studies with transgenic zsGreen-expressing tumors showed that zsGreen was taken up by both cDC1 and cDC2 subsets, as well as monocytes and macrophages (*41*). In our studies, zsGreen was harbored by cDC1 (predominantly migratory), cDC2, MoDC, and pDC in both the superficial and deep cervical lymph nodes, suggesting that multiple DC subsets can traffic antigen from tumor to draining lymph nodes. However, cDC1 are critically required to endogenously prime neoantigen-specific CD8^+^ T cells, and to respond to αPD-L1 therapy against GL261.

We consistently observed that both the superficial and deep cervical lymph nodes contained zsGreen^+^ cDC following intracranial transplantation with CT2A-zsGreen. In previous studies in which Evan’s Blue Dye was injected into the murine ventricles, the deep cervical lymph nodes accumulated blue dye prior to the superficial nodes (*28*). In other work, molecular tracers injected into the brain parenchyma drained only to the deep cervical lymph nodes (*29*). Herein, we observed that a larger percentage of zsGreen^+^ cDC subsets, including cDC1, in zsGreen tumor-bearing mice on day 7 post-transplantation in the deep cervical lymph nodes compared to the superficial cervical lymph nodes. However, in mice examined at 14 days following intracranial tumor injection, the percent of zsGreen^+^ cDC1 was roughly equivalent between the superficial and deep cervical lymph node sites. Although the kinetics of trafficking of molecular dyes from the ventricle to lymph node in Kipnis’ experiments may differ from the trafficking of tumor antigen from brain parenchyma to lymph node, our data support a similar model that deep cervical lymph nodes may sample parenchymal antigens prior to superficial lymph nodes. Ultimately, whereas the drainage route from the venous sinuses to the deep cervical lymph nodes has been well characterized (*28, 29*), the drainage route to the superficial cervical lymph nodes is less well characterized and merits further study.

Parenchymal brain tumor antigens may access the cervical lymph nodes via several potential routes--i.e., via active cell-mediated transit, cellular capture in the lymph nodes of passively drained antigen, or via transport by antigen-bearing exosomes. We used mice lacking CCR7, which is normally required for DC to migrate from the periphery to draining lymph nodes (*59*), to explore whether active cell-mediated migration was required for tumor antigen to be present in the cervical lymph nodes. We consistently observed that a much larger fraction of cDC1 contained zsGreen in the wild-type compared to the CCR7^-/-^ mice, which suggests that antigen trafficking is predominantly an active process that requires intact CCR7-mediated cell migration, which has also been observed in cutaneous melanoma (*41*). However, it is important to note that a zsGreen signal was still observed in DC derived from lymph nodes of CCR7^-/-^ mice, which suggests that either passive drainage mechanisms or cell-mediated transit via other chemokine receptors may contribute to the trafficking of tumor-derived material from the brain to the cervical lymph nodes.

Complementing our findings of zsGreen-containing DC in distinct anatomic locations within and outside brain tumors, we investigated the location of T cell priming in CNS anti-tumor immunity. Given our observations of zsGreen-containing cDC in both the cervical lymph nodes as well as the dura, either location could be a potential site for cDC to present antigen to specific T cells and initiate priming and subsequent clonal expansion. We observed the earliest expansion of OT-I CD8^+^ T cells in cervical lymph nodes, and they consistently harbored the bulk of CFSE-mid, early primed, OT-I CD8^+^ T cells among the tissue sites we examined, which suggests that these were the locations of T cell priming. Moreover, we primarily observed the CFSE-low, terminally divided OT-I CD8^+^ T cells at the later day 6 timepoint, and the majority of them had homed to the tumor. While expanded OT-I cells sometimes appeared in the tumor and dura by the early time point, they were always terminally expanded and CFSE low, which suggests that they had been primed elsewhere. We further observed that both CCR7-deficient and cDC1-deficient mice had delayed, but not absent expansion of adoptively transferred OT-I T cells in mice that harbored GL261-OVA tumors. These observations suggest two things: (1) that tumor antigen can still drain to the cervical lymph nodes in amounts great enough to drive clonal expansion of adoptively transferred T cells despite cells not being carried to lymph nodes by APC, and (2) that cell types other the cDC1 can drive clonal expansion of adoptively transferred T cells if given enough time. However, these data still underscore the importance of cell migration and cDC1 in priming an effective and early T cell response against CNS tumors with normal kinetics. Broadly, our observations extend those of previous work in EAE models in which cervical lymph node resection dampened disease burden (*65–67*), and highlight the importance of the cervical lymph nodes in priming a CNS immune response. More broadly, our data support a model in which tumor antigens from intracranial tumors access the cervical lymph nodes, where subsequent tumor-specific T cell priming can occur.

We coupled our preclinical observations on cDC1 with studies in patients. There has been significant work on the use of DC in GBM immunotherapy. Specifically, DCVax-L is a DC therapy vaccine platform in which autologous monocyte-derived DC are loaded with a patient’s tumor lysate and transferred back into the patient (*79*). Studies have also demonstrated the capacity to enhance the migratory activity of these DC by pre-conditioning to potentially improve patient outcomes (*80*). While these approaches are promising, we focused not on DC as a therapeutic approach in this study, but rather on the location and phagocytic function of endogenously arising cDC subsets in patients undergoing brain tumor surgery. We identified CD141^+^ cDC1, CD1c^+^ cDC2, CD14^+^ monocytes, and CD16^+^ monocytes, as well as CD4^+^ and CD8^+^ T cells in GBM. These findings extend our observations in mice, in which we observed that the tumor is infiltrated with DC far beyond the normal background for steady state brain.

To our knowledge, the phenomenon of direct tumor antigen uptake by cDC has not been observed in human cancers. The use of 5-ALA allows the neurosurgeon to use fluorescence to visually discriminate glioblastoma from normal brain in order to improve the extent of surgical resection (*44–48*). 5-ALA is ingested orally and is selectively metabolized and retained by GBM tumor cells (*48, 49*). We used the transfer of fluorescence principle to probe for the 5-ALA metabolite PPIX in various DC and APC subsets that infiltrated the tumor, which would indicate that the DC had phagocytized part of or a whole tumor cell. Importantly, we observed this phenomenon specifically in tumor-associated APC, as tumor-associated T cells were PPIX^-^. Moreover, cells that acquired and retained PPIX were specifically localized to the tumor, as the identical cell types isolated from PBMCs obtained during surgery after 5-ALA ingestion did not harbor PPIX. We envisage several potential mechanisms of PPIX acquisition by tumor-infiltrating APC: PPIX may be acquired and retained due to the phagocytosis of PPIX^+^ GBM cells, transferred to APC via PPIX-containing GBM exosomes, or other as yet-undefined mechanisms. The selective import of PPIX by CD45^+^ cells appears to be an unlikely mechanism. Interestingly, we observed large fractions (>75%) of PPIX^+^ DC in human GBM, similar to the large fraction (50%-75%) of zsGreen^+^ DC in zsGreen expressing mouse GBM. Moreover, the %PPIX^+^ was not uniform between DC subsets in both humans and mice. These observations have two implications: (1) that different fluorophores with different molecular structures (one a heme-derivative, the other a protein) might be imported into APC by similar mechanisms and (2) that %PPIX^+^ differences observed between differing DC subsets might reflect functional differences between the subsets themselves. Importantly, these findings extend our observations of tumor-antigen uptake in mice to humans.

Growing work has identified the meninges as an immunologically dynamic structure that contains cDC1, cDC2, and pDC in mice (*22, 63*) and in humans (*81*), and that DC within the dura to expand in response to Flt3L as a stimulus (*22, 23*). Recent work has also demonstrated that the dura harbors a network of lymphatic vessels (*28, 29, 31, 55, 68, 82*) that not only drain CSF and antigen to the deep cervical lymph nodes but can proliferate in response to VEGF-C stimulation to help drive a more effective CNS anti-tumor immune response (*55*). Our work extends these findings. We showed that all DC subsets within the dura expanded in response to mice bearing brain tumors, and that all DC subsets, including the cDC1 subset, expanded in response to systemic administration of Flt3L. Furthermore, we identified cDC1 in dura lymphatic vessels by 2 photon microscopy and observed tumor-derived zsGreen within DC harbored by the dura in mice with zsGreen-transduced CT2A brain tumors. Additional studies are needed to define the functional contributions of dura-infiltrating DC, the interplay with the dura lymphatic network, and the anatomic pathways by which parenchymal antigens can access dura immune cell APC. We also translated our preclinical findings to human specimens. We observed, for the first time, that both the CD141^+^ cDC1 human equivalent the CD1c^+^ cDC2 human equivalent infiltrate human dura. Higher resolution studies in the appropriate clinical settings will be needed to further understand their roles in patients in steady state and in CNS disease.

Taken together, these data show that the cDC1 subset is required for immune responses to GBM and infiltrate both tumors and dura mater. Additional work will be needed to understand the anatomic basis of cDC1-associated antigen trafficking to the dura and to the lymph nodes and whether appropriately polarized cDC1 may be leveraged therapeutically in patients with GBM.

## MATERIALS AND METHODS

### Study Design

The objective of this study was to investigate the nature of antigen presentation and the role of cDC1 in CNS anti-tumor immunity. We designed and performed experiments in cellular immunology, and employed preclinical models of GBM, as well as genotypes of mice deficient in cDC1, cell migration, or with lymphatics and/or cDC1 specifically labeled. We used previously characterized neoantigens and model antigens to study antigen-specific responses and kinetics of clonal expansion of adoptively transferred T cells. We also studied APC infiltration and antigen uptake by tumor-infiltrating APCs in patient tumors.

### Mice

All animal experiments were approved by the Washington University Animal Studies Committee. Male and female mice 6-16 weeks of age were used for all experiments. Wild type C57BL/6 mice were purchased from Taconic Biosciences (Hudson, NY). C57BL/6 IRF8+32kb^-/-^ and SNX22^GFP/GFP^ mice were obtained from Dr. Kenneth Murphy. SNX22^GFP/+^ F1 mice were used for experiments. For experiments with GFP-labeled cDC1 and Tomato-labeled lymphatic vessels, we crossed SNX22^GFP/GFP^ C57BL/6 mice to Prox1Cre-ER-tdTomato^+/+^ C57BL/6 mice, obtained from Dr. Gwendalyn Randolph. We treated the F1s as described (*64*) to induce Tomato expression in lymphatic vessels. C57BL/6 CCR7^-/-^ (*83*), OT-I transgenic (*69*), and CD45.1 congenic (*84*) mice were purchased from Jackson. All mice were housed in accordance with IACUC standards.

### Cell lines

CT2A cells were obtained from Dr. Peter Fecci (Duke University) and GL261 cells were obtained from the NCI Tumor Repository. GL261-OFP was generated by transducing parent cells with mOrange2 (obtained from Dr. Robert Schreiber). GL261-zsGreen and CT2A-zsGreen were generated by transducing parent cells with zsGreen (obtained from Dr. David DeNardo). CT2A- mFlt3L was generated by transducing parent cells with murine Flt3L (SinoBiological catalog # MG51113-UT). GL261-OVA was generated by transducing parent cells with ovalbumin cloned from pcDNA3-OVA (Addgene plasmid # 64599). Transduction was performed as described (*85*). All cells were cultured at 37°C, 5% CO_2_ in DMEM with 10% heat-inactivated FBS, 1% Penicillin/Streptomycin, 1% minimum essential amino acids, 1% L-glutamine, and 1% Sodium Pyruvate (D10). Cells were harvested at 70%-90% confluency to inject intracranially.

### Intracranial Injections

50,000 tumor cells were injected 2mm to the right and 2mm posterior of bregma, at a depth of 3.5mm using a Stoelting stereotactic headframe.

### Survival studies

Age and sex matched WT and *IRF8+32kb^-/-^* C57BL/6 mice were intracranially injected with 50,000 GL261 cells as described above. At days 3, 5, 7, and 14, mice were administered by IP injection either PBS vehicle or αPD-L1 (Leinco Technologies, Inc. Clone 10F.9G2) at a dose of 200 µg/mouse in a volume of 100µL. Mice were euthanized when moribund.

### Tissue Harvest

Mice with intracranial tumors were harvested 7-14 days after injection, depending on the experiment. Moreover, for a given study, up to and including the following tissues were harvested: tumor, superficial cervical lymph nodes, deep cervical lymph nodes, a non-draining contralateral inguinal lymph node, dura, and/or spleen. Lymph nodes, dura, and tumor were mechanically dissociated between two frosted slides and digested in 1mg/mL collagenase A, 2%FBS, in RMPI for 20 minutes at 37°C/5% CO_2_. Suspensions were washed and ACK lysed as necessary. Tumors were separated from myelin using a 22.5% Percoll™ solution (Fisher), and centrifuged at RT for 15 minutes at 500g, (acceleration 9, deceleration 5). Mononuclear cells were separated from spleens by first dissociating spleens between a frosted slide, and then using a Ficoll™ gradient (Fisher). We retained the cells in the buffy coat for experiments. In preparation for flow cytometry, cells were suspended in MACS buffer (0.5% BSA, 2mM EDTA in PBS).

## ELISPOT

Tumor single cell suspensions were separated from myelin using a 22.5% Percoll™ (Fisher) solution and subjected to ACK buffer to lyse red blood cells. CD8^+^ T cells were isolated with an EasySep™ Mouse CD8a Positive Selection Kit II (Stem Cell), counted and plated with naïve splenocytes. 50,000 CD8^+^ tumor CD8^+^ T cells were plated with 125,000 naïve splenocytes with or without mImp3 peptide overnight on a pre-coated murine IFN-γ detection plate (Cellular Technologies Limited) and analyzed with an ImmunoSpot plate reader (Cellular Technologies Limited).

### Flow Cytometry and Tetramer

Cell suspensions were filtered, subjected to Fc block, and stained with surface antibodies for >20 minutes on ice. Cells were suspended and flowed in MACS buffer. Flow cytometry was performed on a BD LSRFortessa™ X-20 flow cytometer. Full antibody panel listed in Table S1. The mImp3 (AALLNKLYA)/H-2D^b^ tetramer was generated by the Andrew M. and Jane M. Bursky Center for Immunology and Immunotherapy Programs Immune Monitoring lab as described (*56*). For zsGreen experiments, the 488B/FITC channel was used, with zsGreen^+^ tumor cells as the compensation control. Tumors derived with untransduced tumor cells were used to assess the baseline fluorescence of immune cells isolated from that particular tumor type, and to determine where to draw the positive gate for zsGreen expression by a particular immune cell. Gating definitions described in Table S2.

### 5-ALA Flow Cytometry

5-ALA uptake in U343 cells (obtained from the Cancer Cell Line Encyclopedia (*86*)) was assessed by flow cytometry. U343 cells were incubated with 1 mM 5-Aminolevulinic acid hydrochloride (Sigma) or vehicle as a negative control, trypsinized, and subjected to flow cytometry. In patients who had signed written consent to a Washington University School of Medicine Institutional Review Board-approved protocol, resected tumor and peripheral blood were studied. Tumors were dissociated as described above. PBMC were isolated by centrifuging blood on a Ficoll™ gradient, followed by buffy coat harvest. The 5-ALA metabolite, PPIX, was assessed in the BV650/405C channel and the unstained tumor cell-suspension was used as the compensation control.

### OT-I T cell division assay

CD45.1 mice were crossed to CD45.2/OT-I mice. For adoptive transfer experiments, 5×10^5^ GL261-OVA cells were injected intracranially. Four days later, CD8^+^ T cells were isolated from a CD45.1 x OT1 F1 mouse spleen using an EasySep™ Mouse CD8a Positive Selection Kit II (Stem Cell), CFSE (BioLegend) labeled (10 mins at RT, 5µM), recovered in R10βME at 37°C/5%CO_2_, washed in PBS, and adoptively transferred via tail vein into recipient mice (5×10^5^ OT1 cells per mouse). At days 3 and 6 post adoptive transfer, mice were harvested and tissues dissociated.

### Human tumor and dura preparation

Tumor and dura were obtained from patients who had signed written consent to a Washington University School of Medicine Institutional Review Board-approved protocol. Human dura was macerated and incubated in 2mg/mL collagenase A (Roche) and 2mg/mL collagenase D (Roche), 10% FBS and IMDM overnight at 37°C, with trituration every few hours for dura digestion. The next morning, the single cell suspension was filtered and prepared for flow cytometry.

### 2-Photon Microscopy

Mice were anesthetized and perfused with PBS, the cranial cap was removed with dura still attached to skull, and fixed in ice cold 4%PFA, shaking overnight. If blood vessels were labeled, 5 minutes prior to perfusion, mice were injected IV with 594-lectin (Lycopersicon Esculentum (Tomato) Lectin (LEL, TL), DyLight® 594, Vector Laboratories). For brain sections, brain was cut with a vibrotome after fixation. Fixed tissues were glued to a cover slip with superglue and immersed in PBS for imaging. Images were collected using a custom Leica SP8 two-photon microscope (Leica Microsystems, Wetzlar, Germany) equipped with a 25x 0.95 NA water immersion objective, and two Femtosecond pulsing tunable Ti:Sapphir lasers (Mai Tai HP DeepSee and InSight DS+), both Spectra-Physics (Mountain View, CA, USA). GFP, mOrange and TdTomato were at a wavelength of 925nm whereas Dylight 594 and Dylight 649 were excited at 830nm. Fluorescence emission was guided directly to 4 external detectors in dendritic arrangement (2hybrid and 2 classical PMTs). For signal separation, three dichroic beam splitters (Semrock, Rochester, NY, USA) were used. To separate GFP, mOrange, Dylight 594 and the SHG (Second-harmonic generation) the three cutoff wavelengths were 358nm,538nm and 593nm. The separation of GFP, tdTomato, DyLight 649 and the SHG was obtained with cutoff wavelength of 458, 560 and 652nm. Images were processed and rendered with Imaris cell imaging software.

### Flt3L treatment

Mice were injected subcutaneously in the flank with CT2A-mFlt3L cells, such that the tumor drove expression of Flt3L similar to previously described (*23*). As a negative control, mice were injected with CT2A alone. Mice were harvested 2-3 weeks post-transplant of tumor cells, when flank tumors had reached 1-2cm in diameter.

### Statistical Analysis

Students T-tests were used to analyze differences between groups. Survival statistics were analyzed with log-rank tests. One-tailed T test with Welch’s correction was used to analyze zsGreen data, in which unequal standard deviations were anticipated between groups, and the signal difference between groups was directionally restricted. Unpaired T tests were employed when one mouse was compared to other mice. Paired T tests were used when individual organs within a mouse were compared against one another, or when cell types within an individual patient were simultaneously analyzed and compared. Grubbs outlier test was used when noted. p < 0.05 was considered significant, and statistical analysis were performed with GraphPad Prism 9. Specific statistical test used for each experiment outlined in figure legends.

## Supplementary Materials

Fig. S1. cDC derived from cervical lymph nodes harbor tumor-derived antigen in GL261.

Fig. S2. 2 photon microscopy of dura from *SNX22^GFP/+^* and Prox1-Cre-*TdTomato*^+/WT^ x *SNX22^GFP/+^* mouse under various experimental conditions.

Fig. S3. OT-I expansion in IC CT2A-OVA bearing mouse.

Fig. S4. OT-I expansion in IC GL261 OVA bearing mice is delayed in *IRF8^+^32kb^-/-^* or *CCR7*^-/-^ mice compared to WT mice.

Fig. S5. Fluorescence emission of 5-ALA resected GBM tumor across channels on a conventional BD LSR Fortessa™ X- 20 flow cytometer.

Table S1. Antibody Panel

Table S2. Gating Definitions

## Acknowledgments

We acknowledge the Immune Monitoring Lab (particularly Diane Bender) and Washington University Department of Pathology Flow Cytometry Core for access to flow cytometers and instrumentation. We acknowledge David DeNardo, Gwendalyn Randolph, Josh Rubin, Morey Blinder, and Robert Schreiber for their incisive input and thoughtful contributions. We thank Matthew Holt for his superb illustrations.

## Author contributions

Conceptualization: JABK, GPD

Methodology: JABK, TMJ

Investigation: JABK

Visualization: JABK, BTS, BHZ

Funding acquisition: JABK, GPD

Project administration: JABK, GPD

Supervision: JABK

Writing – original draft: JABK, GPD

Writing – review & editing: JABK, GPD

## Data and materials availability

All data needed to evaluate the conclusions of this paper are present in the paper or in the Supplementary Materials. Any cell lines generated are available upon request.

## References

1. P. Y. Wen et al., Glioblastoma in Adults: A Society for Neuro-Oncology (SNO) and European Society of Neuro-Oncology (EANO) Consensus Review on Current Management and Future Directions. Neuro Oncol, (2020).

2. Q. T. Ostrom et al., CBTRUS Statistical Report: Primary Brain and Other Central Nervous System Tumors Diagnosed in the United States in 2012-2016. Neuro Oncol 21, v1–v100 (2019).

3. R. Stupp et al., Radiotherapy plus concomitant and adjuvant temozolomide for glioblastoma. N Engl J Med 352, 987–996 (2005).

4. G. P. Dunn, H. Okada, Principles of immunology and its nuances in the central nervous system. Neuro Oncol 17 Suppl 7, vii3–vii8 (2015).

5. B. Engelhardt, P. Vajkoczy, R. O. Weller, The movers and shapers in immune privilege of the CNS. Nat Immunol 18, 123–131 (2017).

6. M. Weller et al., Rindopepimut with temozolomide for patients with newly diagnosed, EGFRvIII-expressing glioblastoma (ACT IV): a randomised, double-blind, international phase 3 trial. Lancet Oncol 18, 1373–1385 (2017).

7. R. Maxwell, C. M. Jackson, M. Lim, Clinical Trials Investigating Immune Checkpoint Blockade in Glioblastoma. Curr Treat Options Oncol 18, 51 (2017).

8. T. McGranahan, K. E. Therkelsen, S. Ahmad, S. Nagpal, Current State of Immunotherapy for Treatment of Glioblastoma. Curr Treat Options Oncol 20, 24 (2019).

9. G. P. Dunn, P. E. Fecci, W. T. Curry, Cancer immunoediting in malignant glioma. Neurosurgery 71, 201–222; discussion 222-203 (2012).

10. P. Chongsathidkiet et al., Sequestration of T cells in bone marrow in the setting of glioblastoma and other intracranial tumors. Nat Med 24, 1459–1468 (2018).

11. P. Chongsathidkiet et al., Author Correction: Sequestration of T cells in bone marrow in the setting of glioblastoma and other intracranial tumors. Nat Med 25, 529 (2019).

12. D. A. Wainwright et al., IDO expression in brain tumors increases the recruitment of regulatory T cells and negatively impacts survival. Clin Cancer Res 18, 6110–6121 (2012).

13. L. Zhai et al., Infiltrating T Cells Increase IDO1 Expression in Glioblastoma and Contribute to Decreased Patient Survival. Clin Cancer Res 23, 6650–6660 (2017).

14. E. Ladomersky et al., IDO1 Inhibition Synergizes with Radiation and PD-1 Blockade to Durably Increase Survival Against Advanced Glioblastoma. Clin Cancer Res 24, 2559–2573 (2018).

15. C. W. Brennan et al., The somatic genomic landscape of glioblastoma. Cell 155, 462–477 (2013).

16. E. K. Nduom et al., PD-L1 expression and prognostic impact in glioblastoma. Neuro Oncol 18, 195–205 (2016).

17. A. B. Heimberger, J. H. Sampson, Immunotherapy coming of age: what will it take to make it standard of care for glioblastoma? Neuro Oncol 13, 3–13 (2011).

18. S. M. Razavi et al., Immune Evasion Strategies of Glioblastoma. Front Surg 3, 11 (2016).

19. N. F. Brown, T. J. Carter, D. Ottaviani, P. Mulholland, Harnessing the immune system in glioblastoma. Br J Cancer 119, 1171–1181 (2018).

20. D. J. Aum et al., Molecular and cellular heterogeneity: the hallmark of glioblastoma. Neurosurg Focus 37, E11 (2014).

21. T. Mahlokozera et al., Biological and therapeutic implications of multisector sequencing in newly diagnosed glioblastoma. Neuro Oncol 20, 472–483 (2018).

22. D. Mrdjen et al., High-Dimensional Single-Cell Mapping of Central Nervous System Immune Cells Reveals Distinct Myeloid Subsets in Health, Aging, and Disease. Immunity 48, 599 (2018).

23. N. Anandasabapathy et al., Flt3L controls the development of radiosensitive dendritic cells in the meninges and choroid plexus of the steady-state mouse brain. J Exp Med 208, 1695–1705 (2011).

24. M. K. Matyszak, V. H. Perry, The potential role of dendritic cells in immune-mediated inflammatory diseases in the central nervous system. Neuroscience 74, 599–608 (1996).

25. S. L. Bailey, P. A. Carpentier, E. J. McMahon, W. S. Begolka, S. D. Miller, Innate and adaptive immune responses of the central nervous system. Crit Rev Immunol 26, 149–188 (2006).

26. J. B. Murphy, E. Sturm, Conditions Determining the Transplantability of Tissues in the Brain. J Exp Med 38, 183–197 (1923).

27. A. Waisman, R. S. Liblau, B. Becher, Innate and adaptive immune responses in the CNS. Lancet Neurol 14, 945–955 (2015).

28. A. Louveau et al., Structural and functional features of central nervous system lymphatic vessels. Nature 523, 337–341 (2015).

29. A. Aspelund et al., A dural lymphatic vascular system that drains brain interstitial fluid and macromolecules. J Exp Med 212, 991–999 (2015).

30. S. Da Mesquita, Z. Fu, J. Kipnis, The Meningeal Lymphatic System: A New Player in Neurophysiology. Neuron 100, 375–388 (2018).

31. J. H. Ahn et al., Meningeal lymphatic vessels at the skull base drain cerebrospinal fluid. Nature 572, 62–66 (2019).

32. J. Karman, C. Ling, M. Sandor, Z. Fabry, Initiation of immune responses in brain is promoted by local dendritic cells. J Immunol 173, 2353–2361 (2004).

33. J. Karman et al., Dendritic cells amplify T cell-mediated immune responses in the central nervous system. J Immunol 177, 7750–7760 (2006).

34. A. L. Zozulya et al., Intracerebral dendritic cells critically modulate encephalitogenic versus regulatory immune responses in the CNS. J Neurosci 29, 140–152 (2009).

35. M. G. Harris et al., Immune privilege of the CNS is not the consequence of limited antigen sampling. Sci Rep 4, 4422 (2014).

36. K. Hildner et al., Batf3 deficiency reveals a critical role for CD8alpha+ dendritic cells in cytotoxic T cell immunity. Science 322, 1097–1100 (2008).

37. S. Spranger, R. Bao, T. F. Gajewski, Melanoma-intrinsic beta-catenin signalling prevents anti-tumour immunity. Nature 523, 231–235 (2015).

38. S. Spranger, D. Dai, B. Horton, T. F. Gajewski, Tumor-Residing Batf3 Dendritic Cells Are Required for Effector T Cell Trafficking and Adoptive T Cell Therapy. Cancer Cell 31, 711–723 e714 (2017).

39. H. Salmon et al., Expansion and Activation of CD103(+) Dendritic Cell Progenitors at the Tumor Site Enhances Tumor Responses to Therapeutic PD-L1 and BRAF Inhibition. Immunity 44, 924–938 (2016).

40. M. L. Broz et al., Dissecting the Tumor Myeloid Compartment Reveals Rare Activating Antigen-Presenting Cells Critical for T Cell Immunity. Cancer Cell 26, 938 (2014).

41. E. W. Roberts et al., Critical Role for CD103(+)/CD141(+) Dendritic Cells Bearing CCR7 for Tumor Antigen Trafficking and Priming of T Cell Immunity in Melanoma. Cancer Cell 30, 324–336 (2016).

42. M. B. Fuertes et al., Host type I IFN signals are required for antitumor CD8+ T cell responses through CD8{alpha}+ dendritic cells. J Exp Med 208, 2005–2016 (2011).

43. M. S. Diamond et al., Type I interferon is selectively required by dendritic cells for immune rejection of tumors. J Exp Med 208, 1989–2003 (2011).

44. J. Pastor et al., Role of intraoperative neurophysiological monitoring during fluorescence-guided resection surgery. Acta Neurochir (Wien) 155, 2201–2213 (2013).

45. S. Zhao et al., Intraoperative fluorescence-guided resection of high-grade malignant gliomas using 5-aminolevulinic acid-induced porphyrins: a systematic review and meta-analysis of prospective studies. PLoS One 8, e63682 (2013).

46. Y. Li, R. Rey-Dios, D. W. Roberts, P. A. Valdes, A. A. Cohen-Gadol, Intraoperative fluorescence-guided resection of high-grade gliomas: a comparison of the present techniques and evolution of future strategies. World Neurosurg 82, 175–185 (2014).

47. Z. Huang et al., Fluorescence-guided resection of brain tumor: review of the significance of intraoperative quantification of protoporphyrin IX fluorescence. Neurophotonics 4, 011011 (2017).

48. S. Collaud, A. Juzeniene, J. Moan, N. Lange, On the selectivity of 5-aminolevulinic acid-induced protoporphyrin IX formation. Curr Med Chem Anticancer Agents 4, 301–316 (2004).

49. W. Stummer et al., In vitro and in vivo porphyrin accumulation by C6 glioma cells after exposure to 5-aminolevulinic acid. J Photochem Photobiol B 45, 160–169 (1998).

50. S. Brahler et al., Opposing Roles of Dendritic Cell Subsets in Experimental GN. J Am Soc Nephrol 29, 138–154 (2018).

51. V. Durai et al., Cryptic activation of an Irf8 enhancer governs cDC1 fate specification. Nat Immunol 20, 1161–1173 (2019).

52. C. J. Liu et al., Treatment of an Aggressive Orthotopic Murine Glioblastoma Model with Combination Checkpoint Blockade and a Multivalent Neoantigen Vaccine. Neuro Oncol, (2020).

53. D. A. Reardon et al., Glioblastoma Eradication Following Immune Checkpoint Blockade in an Orthotopic, Immunocompetent Model. Cancer Immunol Res 4, 124–135 (2016).

54. L. Zhai et al., Non-tumor cell IDO1 predominantly contributes to enzyme activity and response to CTLA-4/PD-L1 inhibition in mouse glioblastoma. Brain Behav Immun 62, 24–29 (2017).

55. E. Song et al., VEGF-C-driven lymphatic drainage enables immunosurveillance of brain tumours. Nature 577, 689–694 (2020).

56. T. M. Johanns et al., Endogenous Neoantigen-Specific CD8 T Cells Identified in Two Glioblastoma Models Using a Cancer Immunogenomics Approach. Cancer Immunol Res 4, 1007–1015 (2016).

57. J. Karman, C. Ling, M. Sandor, Z. Fabry, Dendritic cells in the initiation of immune responses against central nervous system-derived antigens. Immunol Lett 92, 107–115 (2004).

58. M. Binnewies et al., Unleashing Type-2 Dendritic Cells to Drive Protective Antitumor CD4(+) T Cell Immunity. Cell 177, 556–571 e516 (2019).

59. R. Forster et al., CCR7 coordinates the primary immune response by establishing functional microenvironments in secondary lymphoid organs. Cell 99, 23–33 (1999).

60. H. B. Stamper, Jr., J. J. Woodruff, Lymphocyte homing into lymph nodes: in vitro demonstration of the selective affinity of recirculating lymphocytes for high-endothelial venules. J Exp Med 144, 828–833 (1976).

61. T. A. Springer, Traffic signals for lymphocyte recirculation and leukocyte emigration: the multistep paradigm. Cell 76, 301–314 (1994).

62. J. P. Lodolce et al., IL-15 receptor maintains lymphoid homeostasis by supporting lymphocyte homing and proliferation. Immunity 9, 669–676 (1998).

63. S. Mundt et al., Conventional DCs sample and present myelin antigens in the healthy CNS and allow parenchymal T cell entry to initiate neuroinflammation. Sci Immunol 4, (2019).

64. R. Bianchi et al., A transgenic Prox1-Cre-tdTomato reporter mouse for lymphatic vessel research. PLoS One 10, e0122976 (2015).

65. G. C. Furtado et al., Swift entry of myelin-specific T lymphocytes into the central nervous system in spontaneous autoimmune encephalomyelitis. J Immunol 181, 4648–4655 (2008).

66. M. J. Phillips, M. Needham, R. O. Weller, Role of cervical lymph nodes in autoimmune encephalomyelitis in the Lewis rat. J Pathol 182, 457–464 (1997).

67. M. van Zwam et al., Surgical excision of CNS-draining lymph nodes reduces relapse severity in chronic-relapsing experimental autoimmune encephalomyelitis. J Pathol 217, 543–551 (2009).

68. A. Louveau et al., CNS lymphatic drainage and neuroinflammation are regulated by meningeal lymphatic vasculature. Nat Neurosci 21, 1380–1391 (2018).

69. K. A. Hogquist et al., T cell receptor antagonist peptides induce positive selection. Cell 76, 17–27 (1994).

70. L. F. Poulin et al., Characterization of human DNGR-1+ BDCA3+ leukocytes as putative equivalents of mouse CD8alpha+ dendritic cells. J Exp Med 207, 1261–1271 (2010).

71. T. L. Murphy et al., Transcriptional Control of Dendritic Cell Development. Annu Rev Immunol 34, 93–119 (2016).

72. S. Hambleton et al., IRF8 mutations and human dendritic-cell immunodeficiency. N Engl J Med 365, 127–138 (2011).

73. J. Yan et al., Author Correction: FGL2 promotes tumor progression in the CNS by suppressing CD103(+) dendritic cell differentiation. Nat Commun 10, 862 (2019).

74. J. Yan et al., FGL2 promotes tumor progression in the CNS by suppressing CD103(+) dendritic cell differentiation. Nat Commun 10, 448 (2019).

75. D. F. Quail, J. A. Joyce, The Microenvironmental Landscape of Brain Tumors. Cancer Cell 31, 326–341 (2017).

76. D. A. Giles, P. C. Duncker, N. M. Wilkinson, J. M. Washnock-Schmid, B. M. Segal, CNS-resident classical DCs play a critical role in CNS autoimmune disease. J Clin Invest 128, 5322–5334 (2018).

77. B. D. Clarkson et al., Innate-adaptive crosstalk: how dendritic cells shape immune responses in the CNS. Adv Exp Med Biol 946, 309–333 (2012).

78. T. Garzon-Muvdi et al., Dendritic cell activation enhances anti-PD-1 mediated immunotherapy against glioblastoma. Oncotarget 9, 20681–20697 (2018).

79. L. M. Liau et al., First results on survival from a large Phase 3 clinical trial of an autologous dendritic cell vaccine in newly diagnosed glioblastoma. J Transl Med 16, 142 (2018).

80. D. A. Mitchell et al., Tetanus toxoid and CCL3 improve dendritic cell vaccines in mice and glioblastoma patients. Nature 519, 366–369 (2015).

81. A. Z. Wang et al., Single Cell Atlas of Human Dura Reveals Cellular Meningeal Landscape and Insights into Meningioma Immune Response. bioRxiv, 2021.2008.2003.454066 (2021).

82. S. Da Mesquita et al., Functional aspects of meningeal lymphatics in ageing and Alzheimer’s disease. Nature 560, 185–191 (2018).

83. U. E. Hopken et al., The chemokine receptor CCR7 controls lymph node-dependent cytotoxic T cell priming in alloimmune responses. Eur J Immunol 34, 461–470 (2004).

84. A. Janowska-Wieczorek et al., Platelet-derived microparticles bind to hematopoietic stem/progenitor cells and enhance their engraftment. Blood 98, 3143–3149 (2001).

85. Y. Fu et al., GATA2 Regulates Constitutive PD-L1 and PD-L2 Expression in Brain Tumors. Sci Rep 10, 9027 (2020).

86. J. Barretina et al., The Cancer Cell Line Encyclopedia enables predictive modelling of anticancer drug sensitivity. Nature 483, 603–607 (2012).

