## Supplementary Material for "The conventional dendritic cell 1 subset primes CD8^+^ T cells and traffics tumor antigen to drive anti-tumor immunity in the brain"

### Supplementary Figures and Tables:

#### Supplemental Figure 1

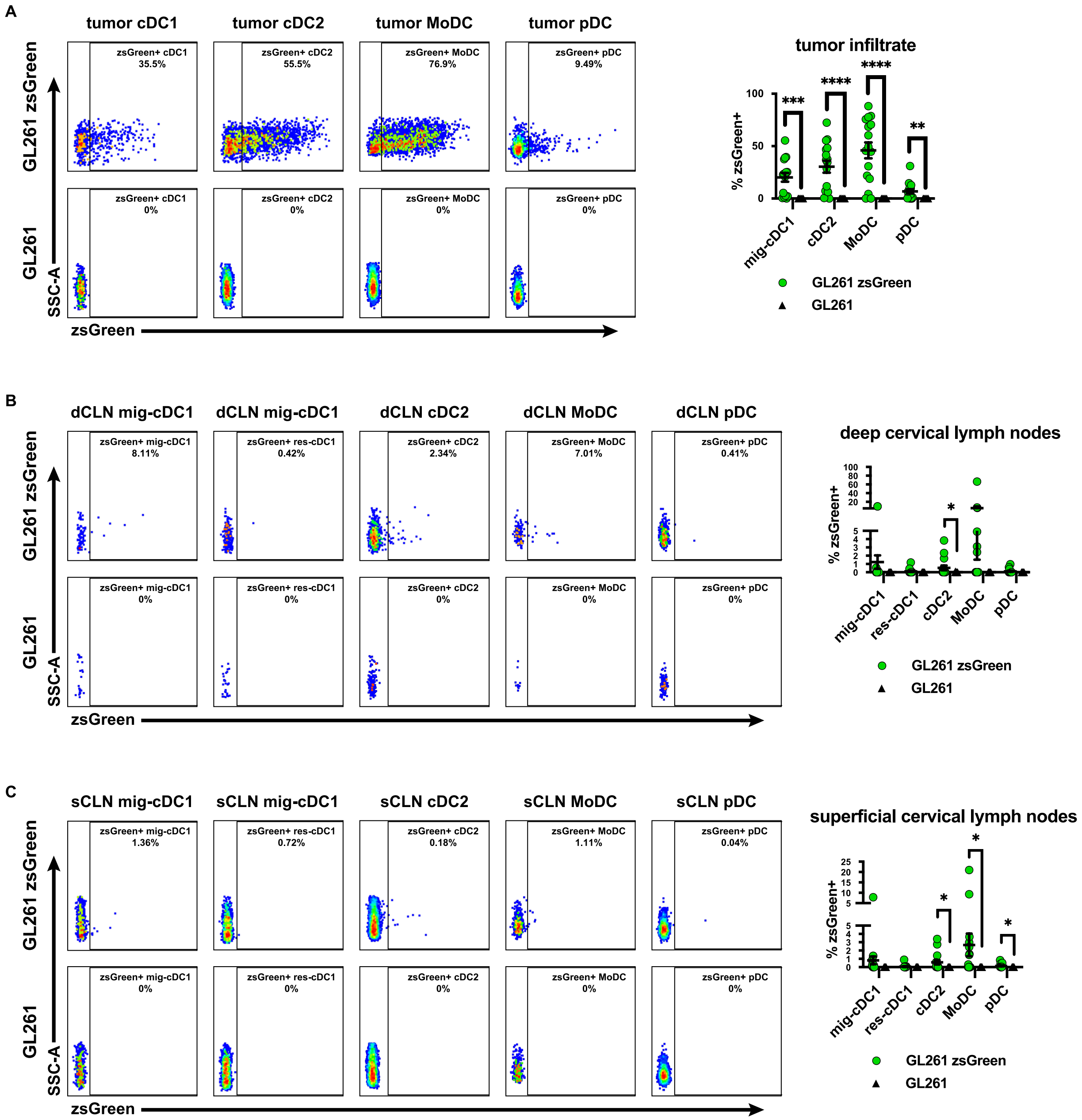

**Supplemental Figure 1. cDC derived from cervical lymph nodes harbor tumor-derived antigen in GL261. (A)** zsGreen uptake by dendritic cells infiltrating GL261-zsGreen tumors. zsGreen expression by **(B)** deep cervical lymph node and **(C)** superficial cervical lymph node dendritic cells. Day 14 tumors for every panel in this figure. Data are represented as mean +/- SEM of at least three independent experiments. *p<0.05, **p<0.01, ***p<0.001, ****p<0.0001. Differences in %zsGreen$^{+}$ analyzed with unpaired one-tailed T test with Welch's correction.

### Supplemental Figure 2

A

-Ctrl

+Flt3L

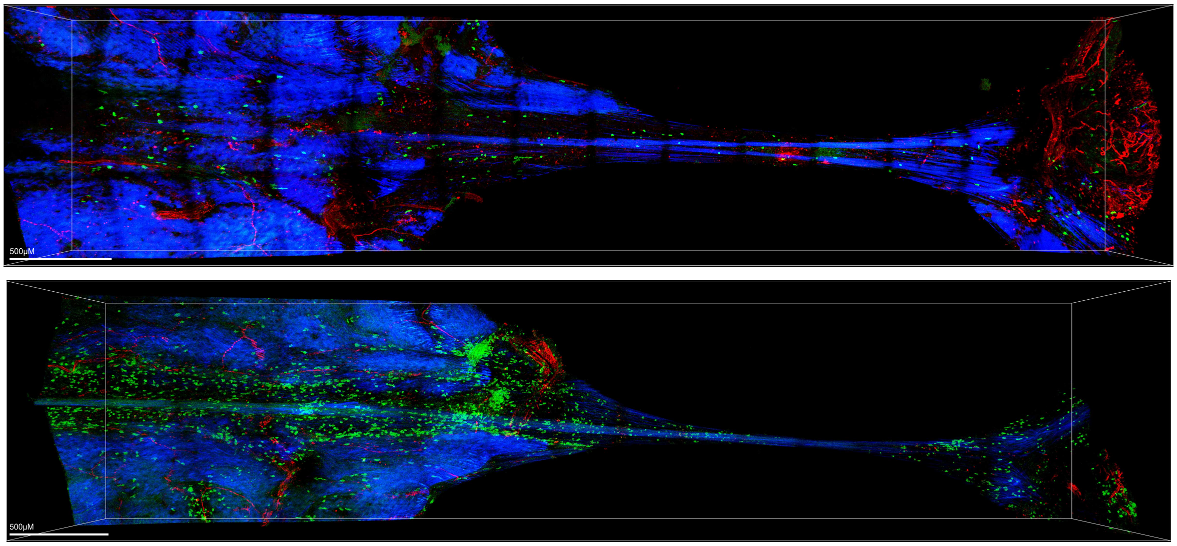


red: lectin (blood vessels) blue: 2nd harmonic (collagen/bone) green: cDC1

B

IC GL261 OFP

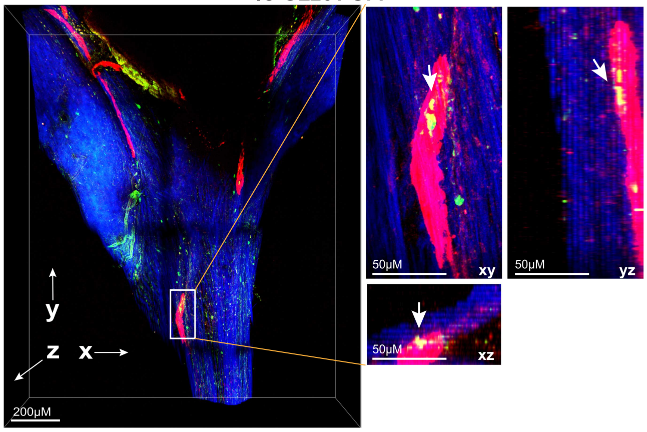


red: Prox1-tomato (lymphatic vessels), blue: 2nd harmonic (collagen/bone), green: cDC1, yellow: cDC1 in lymphatic vessel

C

Sham

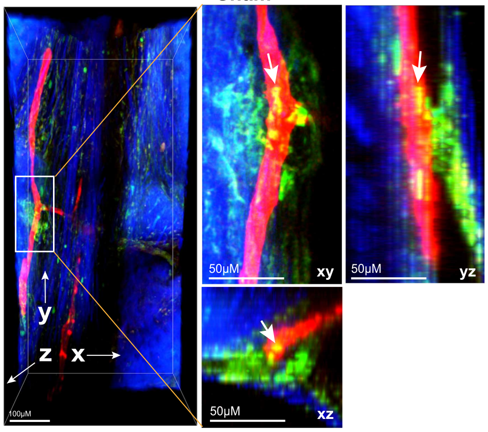


red: Prox1-tomato (lymphatic vessels), blue: 2nd harmonic (collagen/bone), green: cDC1, yellow: cDC1 in lymphatic vessel

**Supplemental Figure 2. 2 photon microscopy of dura from *SNX22*$^{GFP/+}$ and Prox1-Cre-*TdTomato*$^{+/WT}$ x *SNX22*$^{GFP/+}$ mouse under various experimental conditions.** **(A)** 2-photon microscopy of *SNX22*$^{GFP/+}$ mouse dura along the superior sagittal sinus depicting control-treated vs Flt3L treated mice. **(B)** cDC1 harbored within Tomato-labeled lymphatic vessels of IC GL261-OFP injected mouse. **(C)** cDC1 harbored within Tomato-labeled lymphatic vessels IC Sham injected mouse. Dura harvested at day 14 post GL261/Sham injection.

### Supplemental Figure 3

**A**

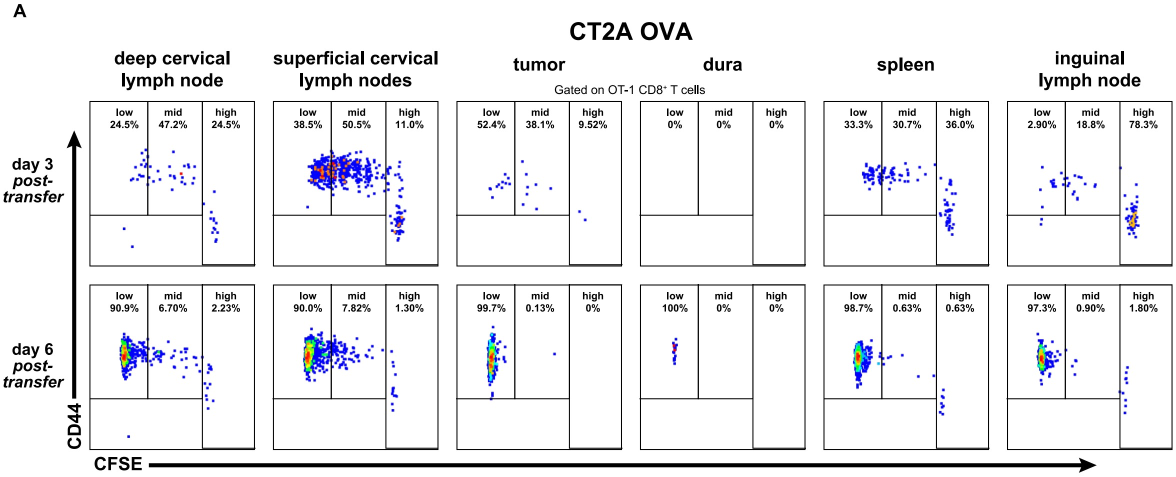

**Supplemental Figure 3. OT-I expansion in CT2A-OVA brain tumor-bearing mouse. (A)** CD44 expression and CFSE dilution of OT-I $CD8^+$ T cells assessed by flow cytometry at day 3 and day 6 post-adoptive transfer of ipsilateral superficial cervical lymph nodes, ipsilateral deep cervical lymph nodes, cerebral hemisphere region encompassing tumor, dura (with tumor abutting region resected), spleen, and non-draining contralateral inguinal lymph node.

### Supplemental Figure 4

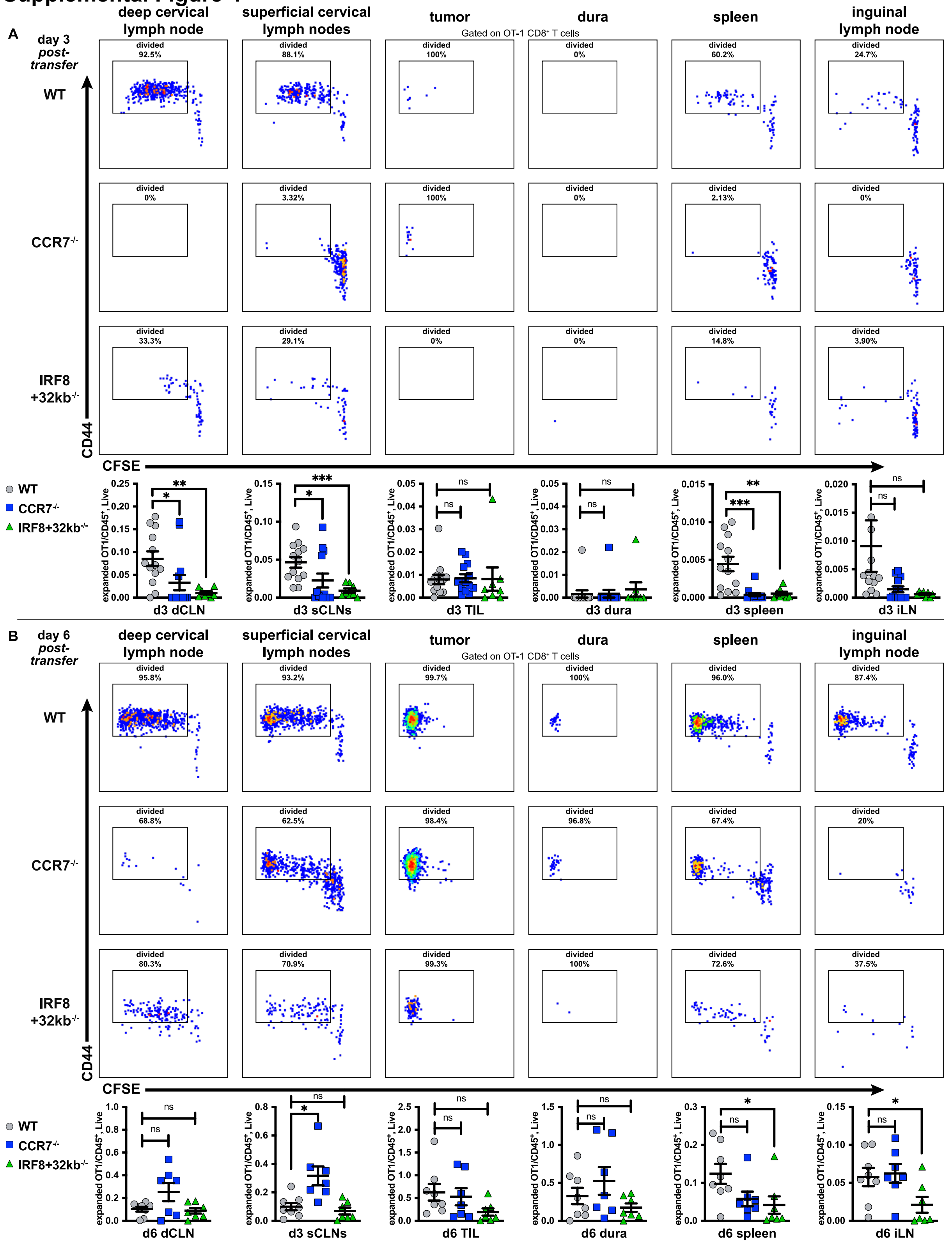

**Supplemental Figure 4. OT-I expansion in IC GL261 OVA bearing mice is delayed in *IRF8*$^{+}$*32kb*$^{-/-}$ or *CCR7*$^{-/-}$ mice compared to WT mice.** CD44 x CFSE dilution of OT-I CD8$^{+}$ T cells at **(A)** day 3 and **(B)** day 6 post-transfer in WT vs. *IRF8*$^{+}$*32kb*$^{-/-}$ or *CCR7*$^{-/-}$ mice, with graphs for each organ below displaying fraction of CFSE-diluted OT-I CD8$^{+}$ T cells over total CD45$^{+}$, live cells. Adoptive transfer was 4 days post intracranial tumor injection. Data are represented as mean +/- SEM of at least three independent experiments. * $p<0.05$, **$p<0.01$, ***$p<0.001$. Data analyzed with unpaired two-tailed T tests.

#### Supplemental Figure 5

**A**

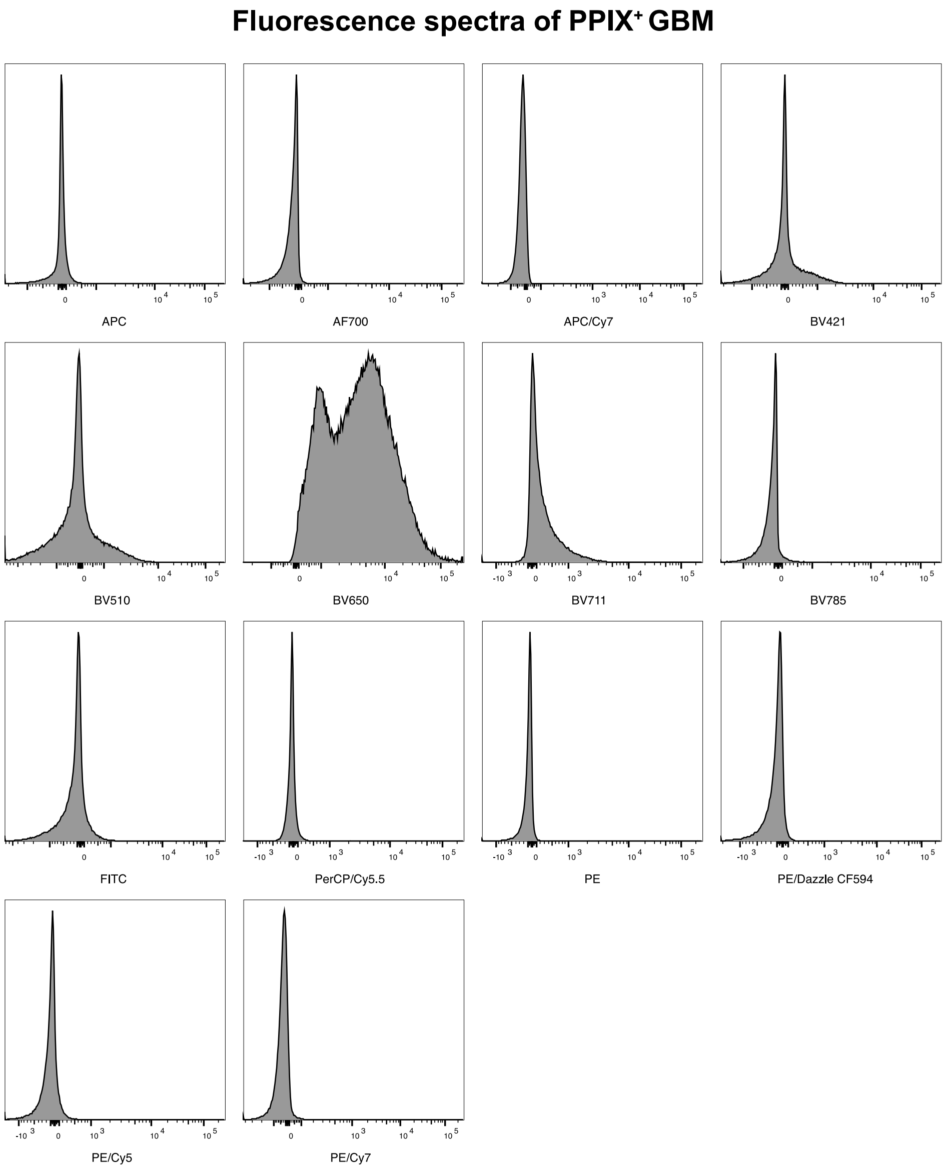

**Supplemental Figure 5. Fluorescence emission spectra of 5-ALA resected GBM tumor across channels on a conventional BD LSR Fortessa™ X- 20 flow cytometer.** All channels used for the experiment are shown.

#### Supplementary Table 1: Antibodies

*Mouse Antibodies*

| Antigen | Color | Vendor | Product no. | Clone |
|---|---|---|---|---|
| **4-1BB** | APC | BioLegend | 106109 | 17B5 |
| **B220** | BV510 | BioLegend | 103248 | RA3-6B2 |
| **CCR7** | PE-Cy7 | BioLegend | 120123 | 4B12 |
| **CD103** | BV421 | BioLegend | 121421 | 2E7 |
| **CD11b** | PE-Cy7 | BioLegend | 101216 | M1/70 |
| **CD11b** | BV650 | BioLegend | 101239 | M1/70 |
| **CD11c** | PE-Cy7 | BioLegend | 117318 | N418 |
| **CD11c** | APC | BioLegend | 117310 | N418 |
| **CD19** | PE-Cy7 | BioLegend | 123113 | 6D5 |
| **CD3** | PE-Cy5 | BioLegend | 100310 | 145-2C11 |
| **CD4** | PerCP Cy5.5 | BioLegend | 100433 | GK1.5 |
| **CD4** | APC | BioLegend | 100412 | GK1.5 |
| **CD44** | BV785 | BioLegend | 103041 | IM7 |
| **CD44** | BV785 | BioLegend | 103041 | IM7 |
| **CD45** | AF700 | BioLegend | 103128 | 30-F11 |
| **CD45** | APC-Cy7 | BioLegend | 103116 | 30-F11 |
| **CD45.1** | BV421 | BioLegend | 110731 | A20 |
| **CD45.2** | AF700 | BioLegend | 109822 | 104 |
| **CD62L** | BV605 | BioLegend | 104441 | MEL-14 |
| **CD62L** | BV510 | BioLegend | 104441 | MEL-14 |
| **CD8a** | PE-Dazzle | BioLegend | 100762 | 53-6.7 |
| **CD8a** | BV711 | BioLegend | 100747 | 53-6.7 |
| **CD8α** | FITC | BioLegend | 100706 | 53-6.7 |
| **CD8α** | PE-Cy7 | BioLegend | 100722 | 53-6.7 |
| **CTLA-4** | BV605 | BioLegend | 106323 | UC10-4B9 |
| **F4/80** | PE-Cy7 | BioLegend | 115519 | BM8 |
| **F4/80** | BV711 | BioLegend | 123147 | BM8 |
| **FoxP3** | PE | BioLegend | 126403 | MF-14 |
| **granzyme B** | AF700 | BioLegend | 372221 | QA16A02 |
| **I-Ab** | AF700 | BioLegend | 107622 | M4/114.15.2 |
| **Ki-67** | FITC | BioLegend | 652409 | 16A8 |
| **Ly-6C** | BV785 | BioLegend | 128041 | HK1.4 |
| **mIFNγ** | BV421 | BioLegend | 505830 | XMG1.2 |
| **NK1.1** | PE-Cy7 | BioLegend | 108713 | PK136 |
| **NK1.1** | BV650 | BioLegend | 108735 | PK136 |
| **OX-40** | PE-CF594 | BioLegend | 119417 | OX-86 |
| **PD-1** | PE-Cy7 | BioLegend | 135215 | 29F.1A12 |
| **PE tetramer** | PE | BioLegend | 405204 | streptavidin |
| **SIRPα** | PE-CF594 | BioLegend | 144016 | P84 |
| **TCR Vα2** | APC | BioLegend | 127809 | B20.1 |
| **TCR Vβ5** | PE | BioLegend | 139503 | MR9-4 |
| **XCR1** | PE | BioLegend | 148204 | ZET |
| **Zombie** | APC-Cy7 | BioLegend | 423106 | n/a |
| **BV421 streptavidin** | BV421 | BioLegend | 405225 | n/a |
| **PE streptavidin** | PE | BioLegend | 405204 | n/a |

*Human Antibodies*

| Antigen | Color | Vendor | Product no. | Clone |
|---|---|---|---|---|
| **CD11c** | PE-Cy5 | BioLegend | 301609 | 3.9 |
| **CD11c** | BV421 | BioLegend | 301627 | 3.9 |
| **CD14** | APC-Cy7 | BioLegend | 367107 | 63D3 |
| **CD14** | BV650 | BioLegend | 301835 | M5E2 |
| **CD141** | APC | BioLegend | 344105 | M80 |
| **CD16** | PerCP-Cy5.5 | BioLegend | 302027 | 3G8 |
| **CD1c** | PE-CF594 | BioLegend | 331531 | L161 |
| **CD3** | PE-Cy7 | BioLegend | 371333 | OKT3 |
| **CD3** | BV510 | BioLegend | 371331 | OKT3 |
| **CD4** | FITC | BioLegend | 357405 | A161A1 |
| **CD45** | AF700 | BioLegend | 368513 | 2D1 |
| **CD8α** | BV711 | BioLegend | 301043 | RPA-T8 |
| **HLA-DR** | BV785 | BioLegend | 307641 | L243 |
| **UV Zombie** | UV | BioLegend | 423107 | n/a |

#### Supplementary Table 2: Gating Definitions

*Mouse*

| Cell type | Definition |
|---|---|
| ***cDC*** | $CD45^{+}$, $F4/80^{-}$, $CD11c^{+}$, $I\text{-}Ab^{+}$, $Ly\text{-}6C^{-}$ |
| ***cDC1*** | cDC plus $XCR1^{+}$, $SIRP\alpha^{-}$ |
| ***migratory cDC1*** | cDC1 plus $CD103^{+}$, $CD8\alpha^{-}$ |
| ***resident cDC1*** | cDC1 plus $CD103^{-}$, $CD8\alpha^{+}$ |
| ***cDC2*** | cDC plus, $XCR1^{-}$, $SIRP\alpha^{+}$ |
| ***pDC*** | $CD45^{+}$, $F4/80^{-}$, $CD11c^{+}$, $I\text{-}Ab^{+}$, $Ly\text{-}6C^{+}$, $CD11b^{-}$ |
| ***MoDC*** | $CD45^{+}$, $F4/80^{-}$, $CD11c^{+}$, $I\text{-}Ab^{+}$, $Ly\text{-}6C^{+}$, $CD11b^{+}$ |
| ***T cells*** | $CD45^{+}$, $CD3^{+}$, $NK1.1^{-}$ |
| ***$CD8^{+}$ T cells*** | T cell plus $CD8\alpha^{+}$, $CD4^{-}$ |
| ***$CD4^{+}$ T cells*** | T cell plus $CD8\alpha^{-}$, $CD4^{+}$; plus FOXP3 for T regulatory cells |
| ***OT-I $CD8^{+}$ T cells*** | $CD45.1^{+}$, $CD45.2^{+}$, $Dump^{-}$ (NK1.1, CD19, CD11b, CD11c, F4/80), $CD3^{+}$, $CD8\alpha^{+}$, $CD4^{-}$, $TCRV\alpha2^{+}$, $TCRV\beta5^{+}$, CFSE variable. |

*Human*

| Cell type | Definition |
|---|---|
| ***cDC*** | $CD45^{+}$, $CD11c^{+}$, $HLA\text{-}DR^{+}$, $CD14^{-}$, $CD16^{-}$ |
| ***cDC1/$CD141^{+}$ cDC*** | cDC plus $CD141^{+}$, $CD1c^{-}$ |
| ***cDC1/$CD1c^{+}$ cDC*** | cDC plus $CD141^{-}$, $CD1c^{+}$ |
| ***$CD14^{+}$ monocytes*** | $CD45^{+}$, $CD11c^{+}$, $HLA\text{-}DR^{+}$, $CD14^{+}$, $CD16^{-}$ |
| ***$CD16^{+}$ monocytes*** | $CD45^{+}$, $CD11c^{+}$, $HLA\text{-}DR^{+}$, $CD14^{-}$, $CD16^{+}$ |
| ***T cells*** | $CD45^{+}$, $CD3^{+}$ |
| ***$CD8^{+}$ T cells*** | $CD45^{+}$, $CD3^{+}$, $CD8\alpha^{+}$, $CD4^{-}$ |
| ***$CD4^{+}$ T cells*** | $CD45^{+}$, $CD3^{+}$, $CD8\alpha^{-}$, $CD4^{+}$ |
